# Predominance of viral core functions over auxiliary metabolism among carbohydrate-active enzymes annotated in the global soil virosphere

**DOI:** 10.1101/2024.12.20.629349

**Authors:** Dominik Merges

## Abstract

Soil viruses are increasingly recognized as components of microbial communities with the potential to alter ecosystem functioning, yet the frequency and functional distribution of virus-encoded metabolic genes in soils remain poorly understood. Here, the Global Soil Virus (GSV) Atlas gene catalog, comprising 1,432,147 viral genes from 1,223 samples across 12 ecosystem types, was searched against KEGG, Pfam, and CAZy for functions related to carbon cycling, nitrogen cycling, and antimicrobial resistance. Annotations associated with host entry or cell-wall lysis were separated from putative auxiliary metabolic functions. Of 5,760 initial gene-level functional annotations across the three categories, 3,902 (67.7%) were classified as peptidoglycanases. Their exclusion left 1,858 genes (0.13% of the catalog), and carbon-cycling functions represented 96.7% of the enzyme-resolved annotations. Chitinase-associated genes formed the largest carbon category, but 78.2% belonged to GH19, a family containing both chitinases and phage endolysins. Among GH19 genes on contigs encoding detectable holins or spanins, 86.8% occurred within three genes of these lysis components, a distribution comparable to that of canonical phage lysozymes. Across six matched metagenomic studies, estimated viral contributions to the targeted functions ranged from 0.10% to 9.86% (mean, 2.60%). Two of the three estimates exceeding 2% were GH19-associated chitinase estimates from rhizosphere soils and were therefore treated as upper estimates. Thus, the putative soil viral metabolic inventory was sparse and concentrated in carbon-degradation functions, whereas much of the initial carbohydrate-active enzyme signal reflected core viral lysis functions rather than auxiliary metabolism.

**IMPORTANCE:** Soil microorganisms drive decomposition and nutrient cycling, and the viruses that infect them may alter these processes by carrying genes that affect host metabolism. However, genes annotated as carbohydrate-degradation enzymes may instead help viruses enter cells or break host cell walls when new virus particles are released. In a worldwide catalog of soil viruses, more than two-thirds of the initial functional assignments represented these infection or release activities. After their removal, putative viral auxiliary metabolic genes were uncommon and concentrated mainly in carbon degradation. Genes potentially involved in chitin degradation were most frequent, but many occurred near viral genes used to break host cells; their estimated contribution should therefore be viewed as an upper limit. Distinguishing core viral functions from host metabolic modification is essential to avoid overestimating viral contributions to soil processes and to identify the strongest candidates for experimental testing.

## INTRODUCTION

Soil viruses are increasingly recognized as abundant components of terrestrial microbial communities and as potential contributors to ecosystem-relevant processes through host lysis and metabolic reprogramming (1, 2). With the compilation of global resources such as the Global Soil Virus (GSV) Atlas, viral genomic potential can now be examined across broad spatial and ecosystem gradients (2). Consequently, metagenomic estimates of community functional potential that are derived from microbial genes alone may be incomplete if virus-encoded functions are not considered.

Distinct responses of bacterial and viral communities to environmental change have been reported across contrasting soil systems. In organochlorine pesticide-contaminated soils, lower bacterial diversity but higher viral diversity and a greater relative abundance of virus-encoded auxiliary metabolic genes (AMGs) linked to pesticide degradation were observed with increasing contamination severity (3). Across an elevational gradient, bacterial metabolic activity was found to decline with increasing elevation, whereas bacteriophage (i.e., viruses that infect bacteria) activity was not significantly affected (4). Active virus–host interactions were also detected in Arctic peat under sub-freezing conditions, where only a small fraction of the detected bacterial community was active (5). Across grassland soils differing in historical precipitation, viral diversity, abundance, lifestyle, and functional potential were found to vary with long-term moisture regime, the driest soils harboring the highest viral diversity and abundance alongside less evidence of virus–host interaction (6). Cohesive viral responses to soil moisture and habitat context have also been reported, suggesting that viral community dynamics can be structured by environmental drivers that do not map one-to-one onto microbial functional profiles (7, 8). Together, these observations motivate an inference risk: functional inventories derived from microbial genes alone may underestimate total gene-centric potential for selected traits and may overattribute those inventories to microbes when viral signals respond differently to environmental drivers.

Interpreting these functional inventories also requires distinguishing shared genomic potential from contributions to ecosystem functioning. Taxonomically distinct microorganisms can encode the same metabolic functions, a pattern commonly discussed in terms of functional redundancy (9). However, such overlap does not establish that taxa are functionally interchangeable across environmental conditions or spatial and temporal scales. Functional similarity has therefore been proposed as a more appropriate term, emphasizing degrees of overlap in specified functions without implying that individual taxa are dispensable (10, 11). In this context, accounting for virus-encoded genes can help resolve how shared functional potential is distributed within metagenomic inventories, while effects on ecosystem functioning require additional evidence.

Auxiliary metabolic genes provide one route through which viruses may contribute to this shared functional potential. AMGs are defined as non-essential, host-derived genes that may modulate host metabolism in ways that increase viral fitness (12). Marine studies have identified AMGs involved in carbon and nutrient transformations (13). Emerson et al. (14) showed that soil viral community composition changed along a permafrost thaw gradient, that some viruses encoded carbon-degradation AMGs such as glycoside hydrolases (GH), and that viral abundances improved prediction of selected carbon chemistry variables beyond host abundance alone. Differences in bacterial genomic potential for carbohydrate utilization and nitrogen transformations have also been detected between managed urban greenspaces and adjacent natural arid soils, while a small set of viral AMGs was recovered in the same comparison (15). Phage-encoded genes associated with carbon metabolism were likewise reported from compost-amended soils (16). However, putative AMG identification is strongly dependent on viral-contig quality, genomic context, functional annotation, and curation, and sequence-based predictions alone cannot establish the biological role or activity of a candidate AMG (12, 17, 18).

Before gene expression can be considered, a more fundamental ambiguity must be resolved: whether a carbohydrate-active enzyme (CAZyme) annotation represents auxiliary metabolism at all. In bacteriophages, glycoside hydrolases and polysaccharide lyases can be carried as virion-associated enzymes that degrade capsular, biofilm, or cell-envelope polysaccharides during adsorption and host entry, while peptidoglycan hydrolases are used during progeny release (19, 20). For canonical phage lysozymes and endolysins, a lytic role is therefore expected rather than an auxiliary metabolic one. In contrast, both chitinases and endolysins are represented within glycoside hydrolase family 19 (GH19), and the relevant substrate—chitin or peptidoglycan—cannot be established from family-level sequence assignment alone (21). Because carbohydrate metabolism has repeatedly been reported as a dominant class of putative soil viral AMGs (2, 22), the extent to which this signal is retained after lysis-associated CAZymes are separated from plausible auxiliary functions remains uncertain. This distinction is therefore evaluated here at the scale of the GSV Atlas before the frequency and ecosystem distribution of virus-encoded metabolic annotations are interpreted.

An effect of a virus-encoded gene on host metabolism can only be produced when the gene is expressed. During productive infection, host metabolism can be transiently redirected by virus-encoded AMGs in ways that favor viral replication (23). For temperate viruses, accessory genes can also be expressed during lysogeny, when viral DNA is maintained as an integrated prophage or, in some cases, as an extrachromosomal phage-plasmid. A host phenotype can thereby be altered by prophage-borne genes, a process termed lysogenic conversion. Phage-encoded virulence factors have long been documented as examples of this process (24), and transfer of antibiotic resistance through lysogenic conversion by functional phage-plasmids has also been experimentally demonstrated (25). Thus, the presence of a gene in a viral genome does not establish whether it is expressed during productive infection or lysogeny, or whether a measurable effect on host metabolism is produced.

Against this background, three functional domains were selected to provide complementary tests of virus-encoded contributions to soil functional potential. Carbon cycling, with emphasis on carbon-degradation functions, was included because carbohydrate-active and other carbon-related genes are among the most frequently reported soil viral AMGs and were a major component of the functional signal identified in the GSV Atlas (2, 22). Nitrogen cycling was included because nitrogen-related viral AMGs have been only sparsely documented in soils, with scattered examples reported rather than evidence for their absence (22). Antibiotic resistance was included as a third, non-nutrient-cycling comparison because phages have been implicated in the dissemination of resistance genes in soils, whereas antibiotic-resistance functions were not separately quantified in the published functional analysis of the GSV Atlas (2). Together, these categories represent a functional domain expected to be comparatively common, one that remains sparsely documented, and one requiring a dedicated annotation framework beyond that used in the original GSV Atlas analysis.

CAZyme annotations were first screened to separate putative auxiliary metabolic functions from virion-structural and lysis-associated functions. The GSV Atlas gene catalog was then used to address three questions: First, how frequently do functionally annotated virus-encoded metabolic genes occur across global soils? Second, are these annotations concentrated in particular functional domains? Third, how large can the virus-encoded component become relative to total metagenomic functional annotations for selected functions within matched viral and microbial datasets? All analyses were restricted to predicted genomic functional potential; gene expression, enzyme activity, and process-level effects were not inferred.

## RESULTS

### Functionally annotated viral metabolic genes were rarely detected across the GSV Atlas gene catalog

Across 1,432,147 viral genes from 1,223 soils, 1,858 genes (0.13%) were assigned functional annotations within the three categories screened here (carbon cycling, nitrogen cycling, and antibiotic resistance) and were supported by at least one of KEGG, Pfam, or CAZy. Of these, 1,169 (0.08% of the GSV Atlas gene catalog) were resolved to a specific enzyme (Table 1; Table S1). All subsequent analyses were based on this enzyme-resolved set; broad, non-enzyme-resolved matches and peptidoglycanase annotations associated with viral lysis were excluded (Table S1). Across ecosystem categories, the enzyme-resolved inventory was dominated by carbon-cycling functions (Fig. 1A). After normalization by total viral gene recovery, broadly similar carbon-cycling annotation rates were observed across the major ecosystem groups. The lowest rate was observed in Peat soils (4.92 per 10,000 viral genes), followed by Unclassified samples (7.41), Agricultural soils (8.03), the pooled Other category (8.39), Grassland (8.99), Permafrost (9.26), and Forest soils (9.50) (Fig. 1A; Table S2). The highest rates were observed in Desert (10.00 per 10,000 viral genes) and Plant-associated soils (10.15). By contrast, nitrogen-cycling annotations were sparsely detected and were restricted to a few ecosystem categories. The largest raw counts were detected in Agricultural (15 genes) and Grassland soils (6 genes), whereas the highest normalized rates were detected in Permafrost (0.51 per 10,000 viral genes), Grassland (0.43), and Agricultural soils (0.42) (Fig. 1A; Table S2). Even fewer antibiotic resistance-associated annotations were detected, with no more than three genes recovered from any ecosystem category and the highest normalized rate detected in Unclassified samples (0.40 per 10,000 viral genes) (Fig. 1A; Table S2). Overall, 1,130 genes (96.7%) were assigned to carbon-cycling functions, whereas only 25 genes (2.1%) and 14 genes (1.2%) were assigned to nitrogen-cycling and antibiotic resistance-associated functions, respectively (Fig. 1B–D; Table 1). Support was provided by a single database for 40.0% of annotations, by two databases for 43.1%, and by all three databases for 16.9% (Table 1).

**Figure 1.**
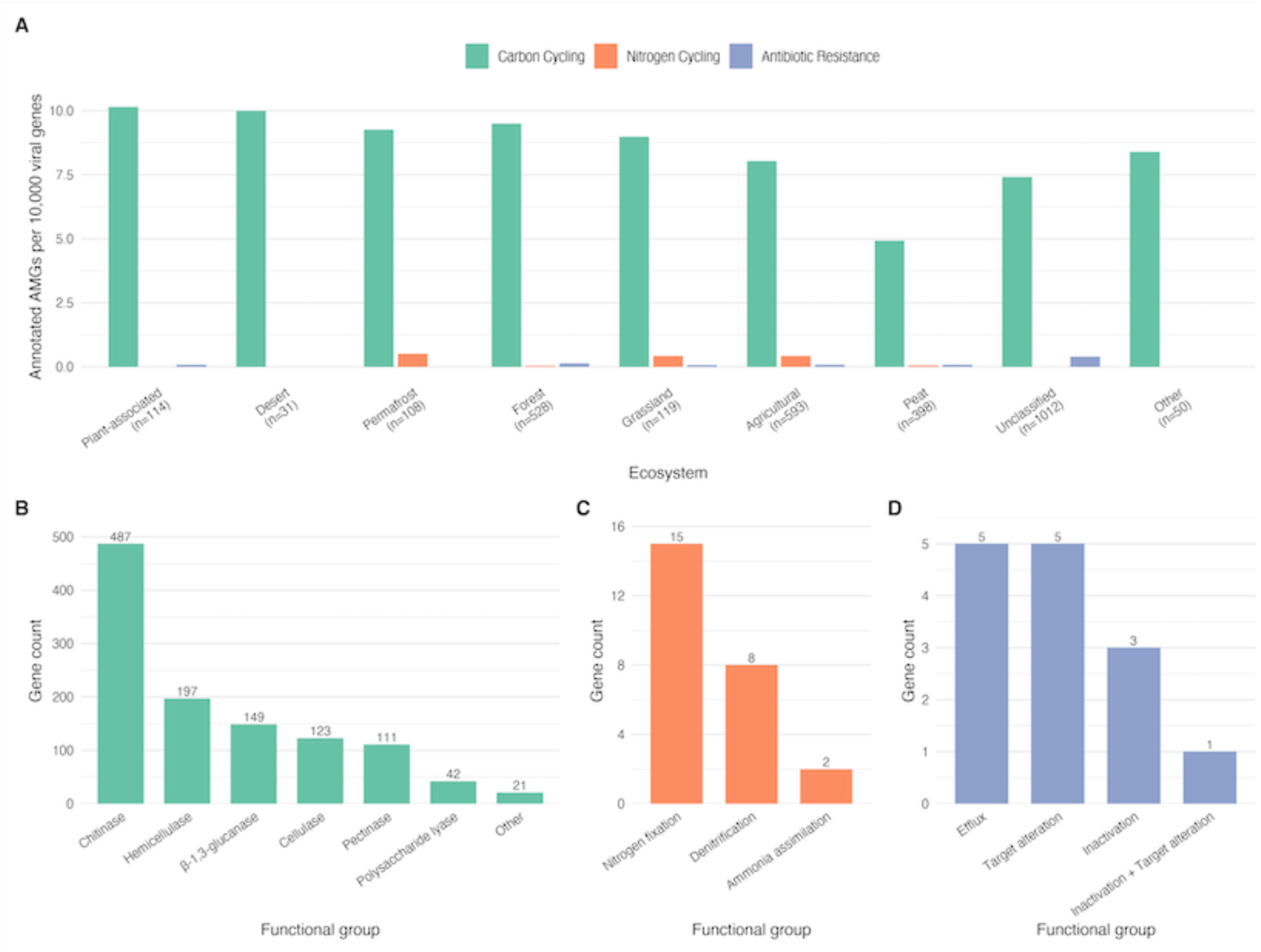
Distribution of putative viral auxiliary metabolic genes (AMGs) across soil ecosystems. (A) Normalized AMG rates (annotated AMGs per 10,000 viral genes) across ecosystem categories, stratified by functional category (carbon cycling, nitrogen cycling, antibiotic resistance). Sample counts per ecosystem are shown in parentheses on the x-axis; these denote all soil samples assigned to each ecosystem in the GSV Atlas metadata (2,953 samples in total). The viral genes analyzed here derive from the 1,223 of these samples that carried at least one recovered viral contig. Ecosystems with fewer than 30 samples were pooled into “Other”; “Unclassified” denotes samples lacking curated ecosystem metadata in the GOLD ontology. (B) Carbon cycling AMGs grouped by functional group. (C) Nitrogen cycling AMGs by functional group. (D) Antibiotic resistance AMGs by functional group. Gene counts are shown above each bar in panels B–D. All panels include only specific (enzyme-resolved) annotations from MetaFuncDecoder v1.0.0; peptidoglycanase (lysis) genes and broad (non-enzyme-resolved) matches were excluded (see Methods).

**Table 1.**
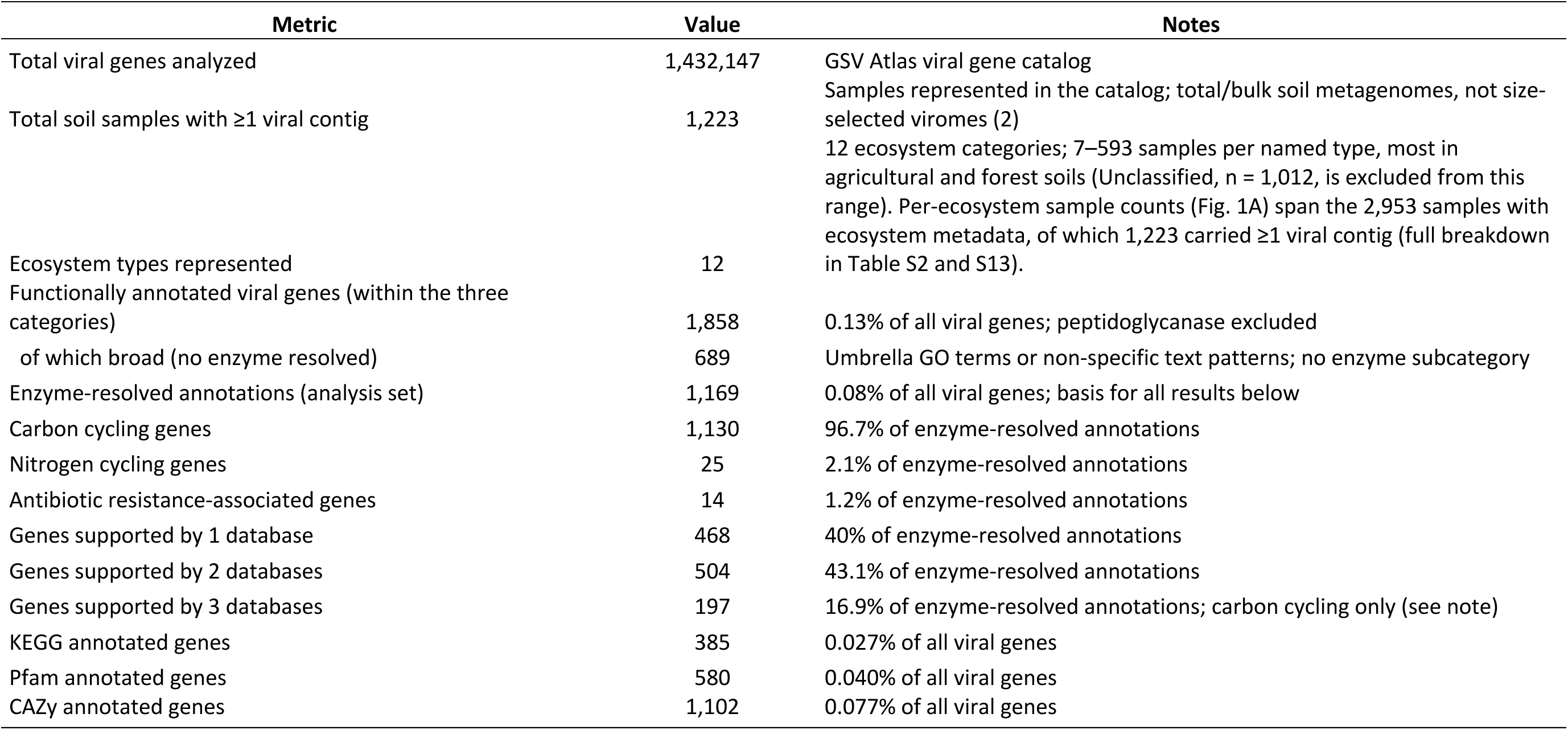
Global soil viral functional landscape overview. All values derived from MetaFuncDecoder v1.0.0. Peptidoglycanase (lysis) genes are excluded throughout; functional and database-support rows refer to the enzyme-resolved analysis set (broad, non-enzyme-resolved matches excluded). The maximum number of supporting databases is 3 for carbon-cycling annotations (CAZy, KEGG, Pfam) but 2 for nitrogen-cycling and AMR annotations (KEGG and Pfam only), because CAZy exclusively indexes carbohydrate-active enzyme families.

### The enzyme-resolved inventory was dominated by carbon-degradation functions, particularly CAZyme-linked activities

Within the carbon-cycling annotations, chitinase-associated functions were detected most frequently (487 genes; 43.1% of carbon-cycling annotations; Fig. 1B; Tables S3 and S4). Of these, 381 genes were assigned to GH19 and 14 to GH18. Because both chitinases and endolysins are represented within GH19, these assignments were retained as chitinase-associated rather than being classified as demonstrated chitin hydrolases. Additional carbon-related annotations were assigned to hemicellulase-associated functions (197 genes; 17.4%), β-1,3-glucanases (149; 13.2%), cellulase-associated functions (123; 10.9%), pectinase-associated functions (111; 9.8%), and polysaccharide lyases (42; 3.7%), with 21 genes (1.9%) assigned to other carbon-related functions (Fig. 1B; Table S3). Among the β-1,3-glucanase annotations, 133 of 149 were assigned to GH16. Agar-, carrageenan-, and porphyran-specific GH16 subfamilies were not detected, and no GH16 annotation was resolved to a marine-algal subfamily. Carbon-cycling annotations were predominantly supported by CAZy, with CAZy support obtained for 1,102 of 1,130 annotations (97.5%; Table S1; Fig. S1). Among CAZy-supported genes, 471 of 1,102 (42.7%) were assigned to chitinase-associated functions, 188 (17.1%) to hemicellulase-associated functions, 149 (13.5%) to β-1,3-glucanases, 123 (11.2%) to cellulase-associated functions, and 42 (3.8%) to polysaccharide lyases (Table S3).

Carbon predominance was similar across the evaluated contig-length classes. Of 49,649 viral contigs, only 80 (0.2%) were assigned to the 1–5 kb size class, and no functional annotations were detected on these contigs. The remaining contigs were distributed between the 5–10 kb (n = 18,225) and ≥10 kb (n = 31,344) size classes. Functional annotations were detected more frequently on ≥10 kb than on 5–10 kb contigs (0.09% versus 0.05%; χ², p < 0.001), whereas similar proportions of carbon-cycling annotations were detected in both size classes (96.7% and 96.1%, respectively; Fisher’s exact OR = 1.20, 95% CI = 0.30–3.47, p = 0.77; Fig. S2).

Within the nitrogen-cycling annotations, 15 of 25 genes (60.0%) were assigned to nitrogen fixation, 8 (32.0%) to denitrification, and 2 (8.0%) to ammonia assimilation (Fig. 1C; Table S3). Nitrogen-fixation annotations were supported primarily by Pfam, whereas denitrification annotations were supported only by KEGG (Table S3). Among the antibiotic resistance-associated annotations, five genes each (35.7%) were assigned to efflux and target alteration, three (21.4%) to antibiotic inactivation, and one (7.1%) jointly to inactivation and target alteration (Fig. 1D; Table S3). Most resistance-associated annotations were supported by a single database, whereas multi-database support was restricted to inactivation-related assignments (Table S3).

### Raw viral CAZyme annotations were dominated by lysis-associated functions

Before lysis-associated annotations were excluded, 5,760 targeted functional annotations were recovered from viral genes. Within the 5,032 enzyme-resolved carbon-cycling annotations, peptidoglycanases represented 77.5% (3,902 of 5,032; Fig. 2A; Table S5), whereas the proportional representation of chitinase-associated annotations would have been reduced from 43.1% in the filtered set to 9.7% in the unfiltered set (Fig. 2A; Table S5).

**Figure 2.**
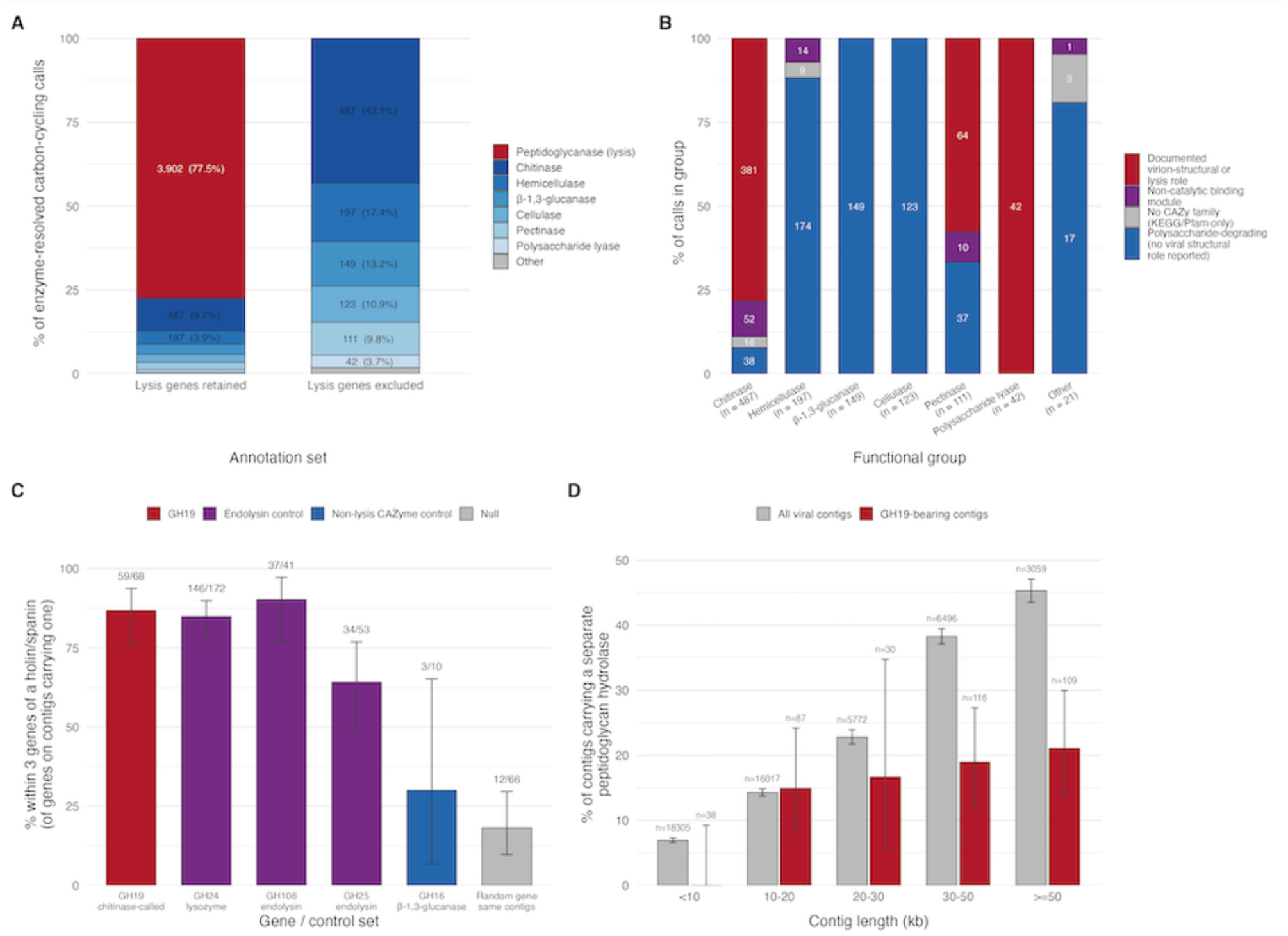
Genes serving core viral functions dominate the annotated CAZyme functions of the soil virosphere. (A) Composition of the enzyme-resolved carbon-cycling annotations with peptidoglycanase (lysis) genes retained (n = 5,032) and excluded (n = 1,130, the analysis set used throughout this study). Counts and percentages are shown for components above 3%. (B) CAZy-family composition of each carbon-cycling group of Fig. 1B classified by whether the family has a documented virion-structural, host-entry or lysis role in viruses. Counts are shown for all segments. (C) Lysis-cassette membership. For genes on contigs that carry a holin or a spanin, the percentage lying within three gene positions of it, with exact binomial 95% confidence intervals; fractions above each bar give the number adjacent out of the number on cassette-bearing contigs. GH24, GH108 and GH25 are endolysin positive controls, GH16 β-1,3-glucanase is a carbohydrate-active enzyme with no reported lysis role, and the null is one randomly drawn non-GH19 gene from each GH19-bearing contig. (D) Percentage of contigs carrying a peptidoglycan hydrolase separate from the query gene, for GH19-bearing contigs and for all viral contigs, stratified by contig length; error bars are binomial 95% confidence intervals and n is the number of contigs per bin. Identifier lists defining peptidoglycan hydrolases, holins and spanins are given in Table S15.

Among the seven retained carbon-function groups, different proportions of annotations were assigned to CAZy families classified as virion-structural or lysis-associated in the present framework (Fig. 2B; Table S6). Cellulase-associated (123/123), hemicellulase-associated (174/197), and β-1,3-glucanase (149/149) annotations were assigned predominantly or exclusively to families not classified as virion-structural or lysis-associated. Among the 487 chitinase-associated genes, 381 (78.2%) were assigned to GH19, 52 (10.7%) to non-catalytic carbohydrate-binding modules, 24 (4.9%) to GH46/GH75 chitosanases, 14 (2.9%) to GH18, and 16 lacked a CAZy family assignment (Table S7). Among the 111 pectinase-associated genes, 64 (57.7%) were assigned to families classified as virion-structural or lysis-associated, and a further 10 were assigned to non-catalytic CBM67 modules (Fig. 2B; Table S6). All 42 polysaccharide-lyase annotations were assigned to alginate-lyase families classified as virion-structural or lysis-associated in the present framework (PL7, n = 33; PL6, n = 5; PL14, n = 4) (Fig. 2B; Table S6).

### Genomic context of viral GH19 genes relative to phage lysis-cassette components

Because GH19 represented the largest retained CAZyme family with ambiguous functional assignment, its genomic position relative to detectable phage lysis-cassette components was examined. Among the 68 GH19 genes located on viral contigs containing a detectable holin or spanin, 59 (86.8%; 95% CI = 76.4–93.8) were located within three ORFs of the detected cassette component, with a median distance of one ORF (Fig. 2C; Table S8). The remaining 313 GH19 genes were located on contigs without a detectable holin or spanin and were therefore excluded from the adjacency analysis. A comparable adjacency frequency was observed for GH24 lysozymes (146/172, 84.9%; OR = 1.17, p = 0.84) and GH108 endolysins (37/41, 90.2%; p = 0.76). A lower adjacency frequency was observed for GH25 endolysins (34/53, 64.2%; OR = 3.62, p = 0.0046), GH16 β-1,3-glucanases (3/10, 30.0%; OR = 14.4, p = 3.6 × 10⁻⁴), and randomly sampled genes from the same contigs (12/66, 18.2%; OR = 28.4, p = 3.7 × 10⁻¹⁶) (Fig. 2C; Tables S8 and S9).

At the contig level, GH19-bearing contigs were stratified by length and were compared with all viral contigs for the presence of a separate peptidoglycan hydrolase (Fig. 2D; Table S10). In the 10–20 kb and 20–30 kb size classes, separate hydrolases were detected on 14.9% versus 14.3% and 16.7% versus 22.8% of GH19-bearing and all viral contigs, respectively. In the 30–50 kb and ≥50 kb size classes, separate hydrolases were detected on 19.0% versus 38.3% and 21.1% versus 45.3% of contigs, respectively.

### Virus-encoded contributions to matched metagenomic functional inventories

To estimate the contribution of virus-encoded genes to broader metagenomic functional inventories, viral-contig-specific annotations were compared with total metagenomic annotations across six matched JGI studies. Five targeted functions were represented: chitinase, ammonia assimilation, nitrogen fixation, and two antibiotic resistance-associated mechanisms (Fig. 3; Tables S11 and S12). Across the six studies, sequencing depths of 25.4–59.4 Gbp were represented (Table S12). For the KEGG-annotated target functions, virus-encoded contributions ranging from 0.10% to 9.86% were estimated, with a mean of 2.60% (Fig. 3B). The largest estimate, 9.86% (7 of 71 genes), was obtained for chitinase, followed by 2.94% (1 of 34 genes) for nitrogen fixation in a different study. A second chitinase comparison yielded an estimate of 2.11% (2 of 95 genes) in another switchgrass-rhizosphere study. In both chitinase-associated comparisons, K03791/PF00182 annotations associated with GH19 were targeted; the resulting percentages should therefore be interpreted as viral representation in this annotated category, rather than as the viral share of experimentally confirmed chitinases. For ammonia assimilation and the two antibiotic resistance-associated mechanisms, estimates of 0.10–0.31% were obtained (Fig. 3B). Across the six exploratory comparisons, viral counts ranged from one to seven and total target-function counts from 34 to 973, and no scaling relationship between viral and total functional-gene counts was detected.

**Figure 3.**
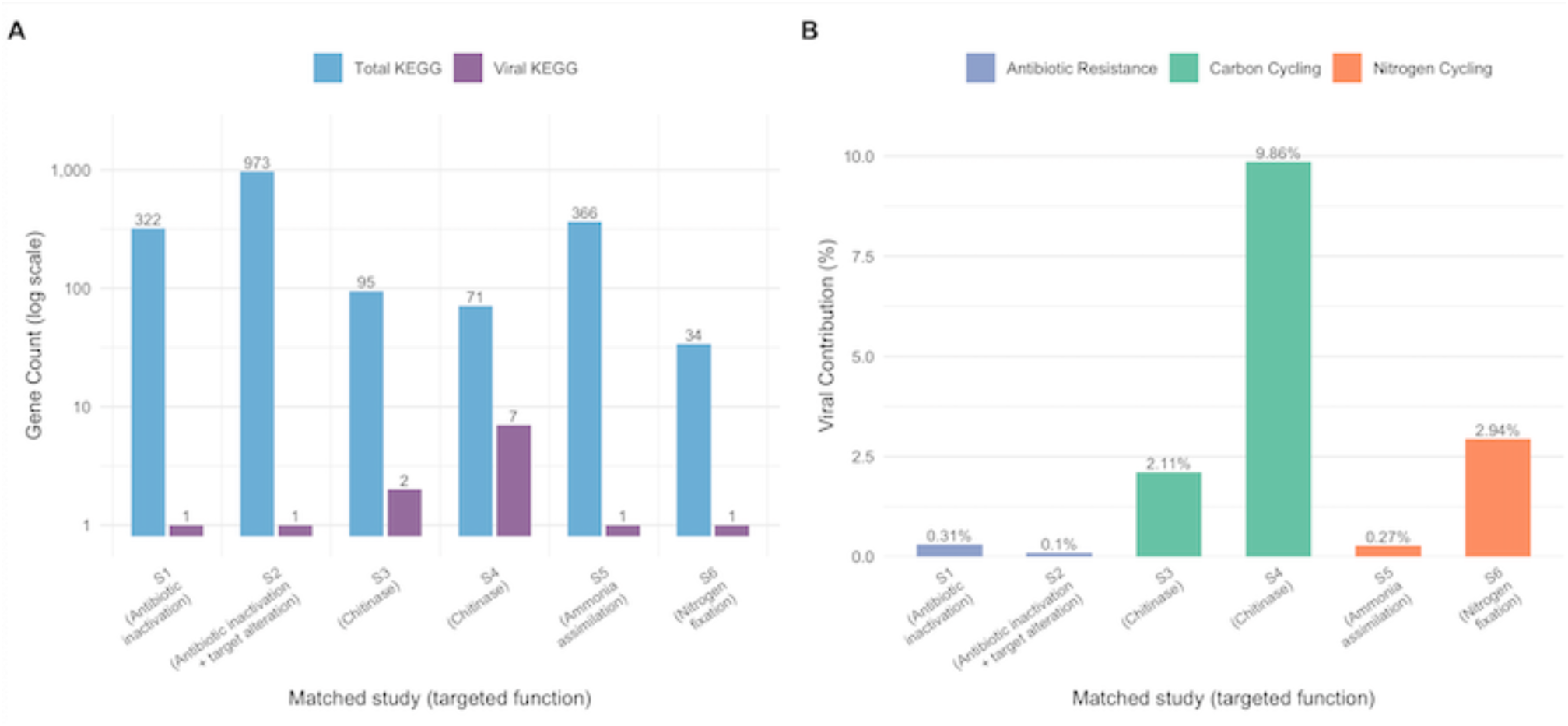
Viral contribution to targeted metagenomic functional annotations in six matched studies. (A) Total and viral KEGG-annotated gene counts (log scale) for five targeted functions: chitinase (carbon cycling), ammonia assimilation and nitrogen fixation (nitrogen cycling), and antibiotic inactivation and antibiotic inactivation + target alteration (antibiotic resistance). (B) Viral contribution as a percentage of total KEGG-annotated genes for each targeted function. The chitinase category of the matched-JGI comparison is K03791/PF00182 in both switchgrass rhizosphere studies, i.e. it is entirely GH19-family and rests on KEGG’s “putative chitinase” orthologue. Functional category labels follow MetaFuncDecoder v1.0.0 subcategory assignments. Full study metadata are reported in Tables S11 and S12.

## DISCUSSION

A quantitative bound on the currently annotatable virus-encoded metabolic potential of global soils was provided by this analysis. Of the 1,432,147 viral genes in the GSV Atlas, no match to KEGG, Pfam, or CAZy was obtained for 1,171,889 genes (81.8%), leaving only 18.2% with a match to at least one of these annotation resources (2). A similarly large fraction of uncharacterized viral sequence space has been reported in previous global virome studies and has been attributed partly to extensive viral sequence divergence and limited representation in reference databases (26, 27). Within the GSV Atlas, only 1,858 genes (0.13% of the complete gene catalog) could be assigned to the three functional domains considered here, and only 1,169 genes (0.08%) were resolved to a specific enzyme. Within this small interpretable fraction, a pronounced functional concentration was detected, with putative carbon-degradation functions accounting for 96.7% of enzyme-resolved annotations and chitinase-associated functions forming the largest individual category. Thus, three features were identified: a large unannotated fraction of the viral gene catalog, a low frequency of annotations within the targeted functional domains, and a strong concentration of the resolved metabolic signal in carbon-degradation functions.

When these findings are placed in the context of microbial biodiversity-ecosystem functioning research, a trait-specific rather than a general contribution of viral genes to ecosystem functional potential is suggested. Carbon-associated annotations remained predominant across contig-length classes and after CAZy-only assignments were removed (Figs. S1–S2). One possible explanation may be provided by differences in the genomic complexity required for individual metabolic functions. Discrete carbohydrate-degradation reactions can be mediated by individual enzymes encoded by single genes, whereas coordinated multigene machinery is required for some nitrogen-cycling functions. Complete nitrogen fixation, for example, requires structural nitrogenase components and proteins involved in FeMo-cofactor biosynthesis (28, 29). Because coding capacity for accessory functions is constrained in compact viral genomes (12), individual catalytic functions may be more readily accommodated than complete multigene pathways. However, this constraint does not necessarily prevent viruses from contributing individual genes to complex host pathways. In the metabolic-reprogramming framework discussed for marine viruses, auxiliary functions can modify existing host pathways in ways that favor viral replication without supplying an entire pathway (23). Whether the soil genes identified here perform such roles remains untested. Exactly one annotated metabolic gene was detected on 1,052 of the 1,108 viral contigs carrying such genes (94.9%), and multiple nitrogen-cycling pathway genes were not detected on any contig. A preference for individual metabolic reactions over complete pathways may therefore contribute to the strong representation of carbon-degradation enzymes in the recoverable viral functional inventory. However, the predominance of single annotated metabolic genes on these contigs does not itself explain the carbon–nitrogen contrast, and pathway complexity alone cannot account for the scarcity of nitrogen-cycling functions.

The scarcity of antibiotic resistance-associated annotations was consistent with previous evidence that antibiotic resistance genes are rarely encoded by phages and can be overestimated by permissive sequence-based annotation. In a survey of 1,181 reference phage genomes, 421 resistance-gene candidates were predicted under exploratory similarity thresholds, whereas only two resistance genes were known; furthermore, antibiotic resistance was not conferred by any of four experimentally tested candidates (30). In a recent comparison of urban greenspace and neighboring natural arid soils, 18 putative viral AMGs were identified, but no antibiotic resistance genes were detected in viral genomes despite a dedicated CARD-based screen (15). The 14 resistance-associated annotations recovered here should therefore be interpreted cautiously. Their low frequency is compatible with genuine scarcity of phage-encoded resistance genes, while sequence-based assignment alone cannot establish a functional resistance phenotype.

The concentration of CAZyme-linked carbon-degradation functions among annotations retained as putative auxiliary metabolic functions was consistent with previous evidence for virus-encoded carbon-related functions in soil viromes. In permafrost peat soils, viral populations carrying putative host-derived metabolic genes were recovered and proposed to influence carbon cycling through metabolic and lysis-mediated effects (31). Virus-encoded glycoside hydrolases with predicted activities against pectin, hemicellulose, starch, and potentially cellulose were subsequently recovered across the same permafrost thaw system, and the activity of a viral GH5 endomannanase was experimentally confirmed (14). More recently, 811 putative viral AMGs were recovered along the same permafrost thaw gradient, nearly half of which were assigned to carbon utilization; these included 59 glycoside hydrolases spanning nine families, several of which were associated with cellulose or hemicellulose degradation (32). These site-specific observations were placed in a broader soil-virome context through analysis of the GSV Atlas. Following activity-specific reclassification, CAZyme-linked carbon-degradation functions were found to remain prominent among annotations retained as putative auxiliary metabolic functions. Nevertheless, the magnitude of the initial carbon-related enzyme signal was substantially reduced when annotations associated with host-cell entry or peptidoglycan degradation during lysis were separated from the putative auxiliary metabolic pool.

The same GSV Atlas gene catalog and CAZy annotation layer as in Graham et al. (2) were used as the starting point for the present analysis, but a different downstream functional screen was applied. The resulting inventory should therefore not be interpreted as a gene-by-gene reassessment of their published putative AMG set. A total of 5,032 carbon-related annotations were recovered, of which 3,902 (77.5%) were assigned to the peptidoglycanase category (Fig. 2A; Table S5). This percentage describes the composition of the inventory screened here and should not be interpreted as a misclassification rate for the original AMG set. In the original GSV Atlas analysis, 3,357 CAZyme-associated genes were reported after putative AMG filtering, and GH73 and CBM50 were among the most frequent CAZy annotations. The broader functional profile was interpreted as indicating potential viral involvement in soil carbon cycling, including organic-matter decomposition and chitin turnover (2). However, the putative status of these assignments and the need for targeted functional assays were also explicitly acknowledged. At the family level, auxiliary carbon metabolism cannot necessarily be distinguished from viral life-cycle functions. GH73 includes N-acetylglucosaminidases involved in bacterial peptidoglycan hydrolysis (33), whereas CBM50 comprises noncatalytic LysM carbohydrate-binding modules through which binding to peptidoglycan or chitin can be mediated (34). Neither catalytic activity nor an auxiliary biological role can therefore be established from a CBM50 assignment alone. The partitioning of a shared annotation resource between likely viral life-cycle functions and candidate auxiliary metabolism can thus be affected by the downstream functional criteria applied.

Greater functional ambiguity was associated with the chitinase-associated signal. Of the 487 chitinase-associated genes, 381 (78.2%) were assigned to GH19, a family in which both chitinase and phage lysis functions have been experimentally demonstrated. Structural similarity between GH19 chitinases and phage lysozymes has long been recognized (35, 36), and contrasting functions have recently been demonstrated for phage-encoded GH19 proteins. A prophage-encoded GH19 chitinase from a deep-sea *Pseudomonas* isolate was shown to support host chitin degradation and growth on chitin (37), whereas a GH19 protein encoded by a cryptic *Pseudomonas aeruginosa* prophage was characterized as a peptidoglycan-active muramidase without detectable chitinase activity (38). A lytic function has also been demonstrated for a cyanophage GH19 protein in the absence of a conventional holin– endolysin system (39). Consequently, GH19 family assignment alone cannot be used to distinguish auxiliary chitin degradation from peptidoglycan hydrolysis.

Additional evidence was provided here by genomic context. Among the 68 GH19 genes located on contigs carrying a detectable holin or spanin, 59 (86.8%) were located within three ORFs of the lysis-cassette component. A comparable adjacency frequency was observed for GH24 lysozymes (84.9%) and GH108 endolysins (90.2%), whereas substantially lower frequencies were observed for GH16 β-1,3-glucanases (30.0%) and randomly sampled genes from the same contigs (18.2%) (Fig. 2C). In the two longest contig classes (30–50 kb and ≥50 kb), a separate peptidoglycan hydrolase was detected approximately twice as frequently on viral contigs overall as on GH19-bearing contigs (Fig. 2D). Taken together, these patterns support a lysis-associated role for a substantial fraction of the GH19 genes for which informative genomic context was available. However, no detectable holin or spanin was present on the contigs carrying the remaining 313 GH19 genes, and their function could therefore not be resolved by the adjacency analysis. Accordingly, the GH19-associated signal should be treated as a set of functionally unresolved candidates, including genes for which a lysis-associated role is supported by genomic context, rather than as a direct measurement of viral chitin-degradation potential.

A smaller component of the carbon-degradation signal was represented by cellulase-, hemicellulase-, and β-1,3-glucanase-associated annotations from families not classified as virion-structural or lysis-associated in the present framework (Fig. 2B). Experimental activity has been demonstrated for a soil viral GH5 endomannanase (14). Separately, a soil viral GH75 protein has been characterized as an active endo-chitosanase (40), providing an experimentally characterized example relevant to the chitosanase-associated component of the present inventory. The plausibility of particular catalytic activities is supported by these characterized proteins; the functions or biological roles of the individual genes recovered here are not thereby validated. Candidates associated with the degradation of structural polysaccharides, including cellulose, hemicellulose, and β-1,3-glucans, can therefore be selected for further investigation on the basis of the functional distinctions applied here, although an auxiliary role cannot be established from these distinctions alone.

At the ecosystem level, a broadly consistent functional profile was detected across the GSV Atlas. After normalization by total viral gene recovery, carbon-cycling annotation rates varied by only approximately twofold across the plotted ecosystem categories (4.9–10.2 genes per 10,000 viral genes), whereas nitrogen-cycling and antibiotic resistance-associated annotations remained rare throughout (Fig. 1A; Table S2). The proportion of viral genes receiving any database annotation was also similar among ecosystem categories, indicating that the large unannotated fraction was not restricted to a particular soil type. This broad functional consistency occurred despite substantial turnover in viral community composition within the GSV Atlas, where few viral operational taxonomic units were shared among geographically separated soils and community composition was associated with environmental gradients (2). Similar environmental structuring of soil viral communities has been reported in relation to moisture, habitat, and soil depth (7, 41). Similar broad annotation profiles were therefore observed across the aggregated ecosystem categories, despite the taxonomic turnover reported for the GSV Atlas.

However, similarity in broad functional composition does not establish that virus-encoded genes contribute equally to total metagenomic functional inventories across soils. To place the GSV-wide patterns in a quantitative metagenomic context, the relative contribution of virus-encoded genes was therefore estimated across six matched datasets. Across these studies, virus-encoded genes represented 0.10–9.86% of the total annotations assigned to the targeted functions, with a mean contribution of 2.60% (Fig. 3). Three of the six estimates exceeded 2%: 9.86% and 2.11% for chitinase-associated annotations in two switchgrass-rhizosphere datasets and 2.94% for nitrogen fixation in peat soil. The two chitinase estimates were derived from K03791/PF00182 annotations corresponding to GH19 and were therefore treated as upper bounds because of the functional ambiguity described above. The nitrogen-fixation estimate, in contrast, was based on a single viral gene among only 34 total K02588 annotations, and was consequently sensitive to the small denominator. Across all six comparisons, between one and seven viral genes were detected against total functional inventories ranging from 34 to 973 genes, and a consistent scaling relationship between viral and total metagenomic gene counts was not detected. Thus, viral representation was strongly dependent on the function and dataset considered rather than being proportional to the overall abundance of the corresponding functional annotation. These findings also inform the functional similarity framework outlined above: virus-encoded genes formed a measurable fraction of annotations assigned to selected functions. Because the viral numerators were small in all comparisons, these estimates should be regarded as exploratory estimates on the contribution of the bulk-metagenome-recoverable viral fraction rather than as precise ecosystem-level effect sizes.

The quantitative estimates obtained here should therefore be interpreted as conditional on both viral recovery and functional classification. Because the GSV Atlas and the matched comparisons were derived from bulk metagenomes, only the viral fraction recoverable by total-community sequencing was represented. Substantially different viral populations can be recovered from the same soil by bulk metagenomes and size-selected viromes, with greater recovery of rare viral populations generally obtained by virus-targeted sequencing (42). The percentages reported here should therefore be interpreted as estimates for the bulk-metagenome-recoverable viral fraction rather than for the complete soil virosphere. Putative AMG inventories are also shaped by upstream viral identification and downstream functional curation. In DRAM-v, genomic-context scores and metabolic flags are combined, and genes associated with viral attachment or entry, including many CAZymes, are excluded from default auxiliary assignments (43), whereas candidate auxiliary functions are identified through a distinct annotation and curation framework in VIBRANT (44). In the GSV Atlas, putative AMG prediction excluded proteins classified by geNomad as virus-specific and applied genomic-context and functional-annotation filters (2). Although this procedure could remove recognized viral housekeeping proteins, an explicit CAZyme-family-level assessment of potential lysis functions was not described. Even after candidate genes have been recovered, their ecological interpretation is further affected by the functional resolution of the underlying databases and by how individual families are assigned to metabolic or viral processes. Downstream comparability can be improved by the lysis-aware family-to-function mapping implemented in MetaFuncDecoder v1.0.0 (45), but differences in upstream viral recovery or AMG detection cannot be eliminated by this approach, and classification remains dependent on the accuracy and resolution of sequence-based annotations (12). Both the recovered viral sequence space and the criteria used to define candidate auxiliary functions should therefore be considered in comparisons across studies.

More fundamentally, the present annotations establish gene presence rather than gene activity. Even when a functional assignment is correct, metabolic potential cannot be equated with transcription, protein production, enzymatic activity, or a measurable effect on host or ecosystem processes. None of the soil viral GH19 genes or other candidate auxiliary metabolic genes identified here was evaluated at the transcript or protein level, and it therefore remains unknown whether these genes are expressed during infection or lysogeny and whether the encoded proteins are enzymatically active. The same limitation applies to the few antibiotic resistance-associated genes detected, for which sequence-based annotation alone cannot establish a resistance phenotype or a contribution to horizontal gene transfer. Demonstration of a physiological role will therefore require complementary metatranscriptomic, proteomic, or biochemical evidence. Finally, the present analysis was restricted to carbon cycling, nitrogen cycling, and antibiotic resistance. Other virus-encoded functions implicated in soil biogeochemistry and host physiology, including phosphorus and sulfur cycling, metal detoxification, pollutant degradation, transport, and energy metabolism, were not considered here (22). Whether the trait-specific pattern identified in the present analysis extends to these additional functional domains therefore remains unresolved.

## CONCLUSION

By applying a lysis-aware functional classification, a substantial fraction of the apparent CAZyme signal in the soil virosphere was resolved as being associated with core viral functions rather than auxiliary metabolism. After these functions were separated, virus-encoded metabolic potential remained strongly concentrated in carbon-degradation activities, but the confidence of this interpretation differed among enzyme families. In particular, GH19-associated chitin degradation remained functionally ambiguous, whereas cellulase-, hemicellulase-, and β-1,3-glucanase-associated genes represented comparatively well-supported candidates for auxiliary carbon-degradation functions. In matched metagenomes, virus-encoded contributions of up to 9.86% were estimated for selected functions, although the largest value represented a GH19-based upper estimates. Excluding virus-encoded genes from function-specific metagenomic inventories may therefore omit a measurable component of community genomic potential. The present estimates remain restricted to genomic potential, and metatranscriptomic, proteomic, and biochemical validation will be required before effects on host metabolism or ecosystem carbon turnover can be inferred. Overall, the results indicate that virus-encoded carbon-degradation potential is detectable across soils, but that its magnitude can be substantially overestimated when CAZyme annotation is interpreted without considering viral lysis functions.

## MATERIALS AND METHODS

### Global analysis of virus-encoded functional annotations

The GSV Atlas published by Graham et al. (2) was reanalyzed, with a focus on the gene catalog associated with soil viral contigs. Viral sequences from a global collection of soil metagenomes are compiled in the GSV Atlas, from which 1,432,147 viral genes were recovered. The viral gene annotation tables provided with that resource were used to characterize the functional landscape of soil viral communities across 1,223 soil samples spanning 12 ecosystem types. All genes analyzed here were derived from assembly-based annotation within the GSV Atlas pipeline (2). In brief, 1,552 metagenomes not previously included in IMG/M were analyzed with the JGI Metagenome Workflow and individually assembled with metaSPAdes v3.1 (46, 47), whereas 1,480 curated samples were obtained from IMG/M with existing assemblies. Resulting new assemblies were processed through the IMG/M Metagenome Annotation Pipeline v5.0.0 (48). Gene calls and associated functional annotations were taken directly from the GSV Atlas/IMG framework and were not recomputed here (2). Viral contigs were identified with a modified IMG/VR v3 pipeline supplemented by geNomad v1.3.3 and were quality-assessed with CheckV v1.0.1 (2, 49, 50), yielding 49,649 quality-controlled viral contigs carrying 1,432,147 gene calls. No read-based gene detection or read-mapping abundance estimation was performed; only unique gene calls on distinct assembled contigs were counted, with each call included once regardless of sequencing depth and without coverage-based weighting.

### Functional annotation and category assignment

Functional category assignment (carbon cycling, nitrogen cycling, and antibiotic resistance) was performed with MetaFuncDecoder v1.0.0 (45), a Python tool developed for this study. The pre-existing per-gene KEGG, Pfam, and CAZy annotations distributed with the GSV Atlas (2) were read directly, and viral genes were not reannotated. Hits from each database were matched independently against activity-specific rules. CAZy families were matched through an explicit family-to-activity list based on López-Mondéjar et al. (51) and the CAZy family activity descriptions of Drula et al. (52); KEGG orthologs were matched through text patterns naming the same activity; and Pfam domains were matched through their mapped Gene Ontology terms (53). Nitrogen-cycling patterns were based on NCycDB (54), and antibiotic-resistance patterns were based on the CARD Antibiotic Resistance Ontology (55). A subcategory was assigned when a match to the corresponding rule was obtained for at least one available hit, rather than by cross-referencing identifiers across databases. Within carbon cycling, fine-grained enzyme subcategories were resolved by MetaFuncDecoder (for example, chitinase, cellulase, β-glucosidase, mannanase, xylanase, arabinofuranosidase, β-1,3-glucanase, pectinase, alginate/polysaccharide lyase, starch degradation, and oxidative/lignin-active enzymes); for reporting, these were collapsed into seven carbon-activity groups: chitinase, hemicellulase, β-1,3-glucanase, cellulase, pectinase, polysaccharide lyase, and a residual “other” group (Fig. 1B). Nitrogen-cycling annotations were classified as nitrogen fixation, denitrification, or ammonia assimilation, and antibiotic-resistance annotations were classified as efflux, target alteration, or antibiotic inactivation.

Each gene was additionally assigned a database-support score (one, two, or three) according to the number of databases in which the same subcategory label was independently obtained. Because only carbohydrate-active enzyme families are indexed by CAZy, CAZy support was unavailable for nitrogen-cycling or antibiotic-resistance functions; the maximum attainable support was therefore three for carbon-cycling annotations but two for nitrogen-cycling and antibiotic-resistance annotations (KEGG and Pfam only). Database-support counts are reported within each functional category so that this structural difference is not conflated with annotation quality across categories. A subcategory was assigned by MetaFuncDecoder only when a specific enzyme was resolved; no subcategory was assigned to matches supported solely by umbrella Gene Ontology terms or nonspecific text patterns, which were termed broad. Main analyses used specific assignments; Fig. S1 and Table S14 additionally summarize broad assignments. Figure S1B compares three alternative filters; its KEGG/Pfam-supported set includes broad matches.

### Exclusion of peptidoglycanase-associated annotations

To reduce the inclusion of likely viral life-cycle functions in the targeted functional inventory, MetaFuncDecoder v1.0.0 was used to identify and exclude peptidoglycanase-associated annotations. Retained annotations were subsequently assessed for additional functional ambiguity. A dedicated peptidoglycanase subcategory was assigned to the CAZy families GH22, GH23, GH24, GH25, GH73, GH102, GH103, GH104, GH108, GH153, and CE9, following the “Peptidoglycanases” classification of López-Mondéjar et al. (51), and to KEGG orthologs and Pfam domains describing a lysozyme, muramidase, N-acetylmuramidase, peptidoglycan hydrolase, or lytic transglycosylase. These genes were excluded from the main enzyme-resolved metabolic inventory unless explicitly retained for the filter-comparison analysis. Related curation principles are reflected in the DRAM-v attachment (A) flag, through which genes associated with viral host attachment and entry, including many CAZymes, are marked for exclusion from default AMG summaries (43). A complementary example is provided by Luo et al. (56), in whose manual AMG curation glycoside hydrolases and peptidases associated with cell-wall lysis were excluded. Here, the peptidoglycanase category was assigned directly by MetaFuncDecoder from the existing annotations, allowing a separate screen to be performed without DRAM-v output, including the GSV Atlas.

### Effect of the peptidoglycanase filter on inventory composition

To quantify the effect of lysis-associated annotations on the apparent carbon signal, the composition of the enzyme-resolved carbon-cycling annotations was compared with and without the peptidoglycanase subcategory. Annotations were grouped into the seven carbon-activity classes used throughout (chitinase, hemicellulase, β-1,3-glucanase, cellulase, pectinase, polysaccharide lyase, and other), with peptidoglycanase forming an eighth group when retained. Each group was expressed as a percentage of the total under each scenario: all enzyme-resolved carbon-cycling annotations and the same set after removal of the peptidoglycanase group.

### Classification of retained carbon-degradation families

After exclusion of the peptidoglycanase subcategory, each remaining enzyme-resolved carbon-cycling annotation was assigned to one of four mutually exclusive classes on the basis of its CAZy family to separate annotations interpretable as putative carbon-degradation functions from those attributable to core viral functions. A “no CAZy family” class was used for annotations resolved only by a KEGG ortholog or a Pfam domain, with no CAZy family assignment. Among annotations carrying a CAZy family, noncatalytic carbohydrate-binding modules, which were treated as providing substrate binding but no hydrolytic activity, were classified separately. Families for which a documented virion-structural, host-entry, or lysis role has been reported in viruses—the chitinase/endolysin family GH19, the polygalacturonase GH28, and six pectin and alginate lyase families—were assigned to a third class (19, 21, 38, 39). All remaining families, for which degradation of plant-, fungal-, or bacterial-derived polysaccharides has been reported but no viral structural role has been described, were assigned to the fourth class. Full identifier sets for all classes are listed in Table S15.

### Genomic context of GH19 annotations

To assess whether GH19 genes assigned to the chitinase category were located in genomic positions characteristic of a phage endolysin, every gene in the catalog was flagged for the three components of the phage lysis cassette. Peptidoglycan hydrolases were identified across all endolysin catalytic classes (57, 58): the glycosidase class—eleven CAZy families, six glycoside-hydrolase Pfam domains, and nine KEGG orthologs—together with the amidase and endopeptidase classes, which were captured by eight additional Pfam domains (Table S15). In this broader genomic-context scan, peptide- and amide-bond-cleaving endolysins were therefore recovered in addition to glycosidase-class endolysins. Holins were identified from 36 Pfam families for which a holin or antiholin designation was provided in the entry name, and spanins were identified from three Pfam families (full identifier lists in Table S15). These assignments were taken from the per-gene Pfam, CAZy, and KEGG annotations distributed with the GSV Atlas and were not recomputed. Within each contig, genes were ordered by start coordinate, and the distance, in gene positions, from every query gene to the nearest holin or spanin on the same contig was determined; a gene was scored as cassette-associated when this distance was three genes or fewer. GH19 chitinase-assigned genes were tested against three endolysin families as positive controls (GH24 phage lysozyme, GH108, and GH25), against GH16 β-1,3-glucanase as a carbohydrate-active enzyme with no reported lysis role, and against a null set for which one non-GH19 gene was randomly selected from each GH19-bearing contig under a fixed random seed. Because a cassette marker can only be located on contigs where a holin or spanin was annotated, the adjacency test was restricted to genes residing on such contigs; the proportion of cassette-associated genes was compared between GH19 and each control set by two-sided Fisher’s exact test and was reported with exact binomial (Clopper-Pearson) confidence intervals. To account for the greater probability that a second lysis gene would be recovered on longer contigs, the carriage of an additional peptidoglycan hydrolase was compared between GH19-bearing contigs and all viral contigs. Each contig was scored as positive if at least one peptidoglycan hydrolase, drawn from the same CAZy, KEGG, and Pfam identifier set and excluding the focal GH19 gene, was detected; contigs were stratified into five length classes (<10, 10– 20, 20–30, 30–50, and ≥50 kb), and the proportion of positive contigs in each class was compared between the two sets with exact binomial confidence intervals.

### Ecosystem classification and normalization

Ecosystem labels were assigned to each metagenome from the GOLD ontology fields Ecosystem Type and Specific Ecosystem as curated in IMG/M. Synonymous or inconsistently capitalized labels were harmonized (for example, “Forest Soil” and “Forest soil” were merged as “Forest”; “Agricultural land”, “Agricultural”, and “Agricultural soil” were merged as “Agricultural”; full mapping in Table S16), and plant-associated types (Rhizosphere, Roots, and Rhizoplane) were combined into a single category. Samples without specific ecosystem metadata were classified as “Unclassified”, and ecosystems represented by fewer than 30 samples were pooled as “Other”. To account for differences in sampling effort and viral gene recovery across ecosystems, annotation counts were normalized as the number of annotated genes per 10,000 viral genes per ecosystem. For each ecosystem, the proportion of viral genes matching at least one of Pfam, KEGG, or CAZy is additionally reported in Table S13 alongside the category-specific rate.

### Contig-length stratification

To test whether the enrichment of carbohydrate-active annotations was driven by short, potentially unreliable contigs, all functional results were stratified into three contig-length bins (1–5 kb, 5–10 kb, and ≥10 kb). The ≥10-kb threshold was selected because the GSV Atlas applied different viral-identification criteria above and below 10 kb, consistent with MIUViG guidance that short (<10 kb) viral sequences should be interpreted cautiously (2, 59). Peptidoglycanase (lysis) genes were excluded before stratification, and the proportion of carbon-cycling annotations was compared between the 5–10-kb and ≥10-kb bins by Fisher’s exact test.

### Comparative analysis of viral and total metagenomic functional annotations

To estimate the potential effect of virus-encoded functions on gene-centric functional inventories, viral annotations were compared with total metagenomic annotations in matched soil datasets retrieved from the JGI Integrated Microbial Genomes and Microbiomes (IMG/M) platform on 10 January 2026. Study identifiers linking GSV Atlas entries to their source metagenomes were provided by Graham et al. (2), so that exact matching between the viral datasets and the corresponding total metagenomes was possible. Six matched studies (Tables S11 and S12) were selected for a exploratory comparison spanning carbon-degradation functions (two agricultural-rhizosphere studies), nitrogen-related functions (two peat studies), and antibiotic-resistance functions (agricultural-rhizosphere and forest studies). In every case, viral contigs were identified within the same assembled metagenome from which the total annotations were derived; viral and total annotations were therefore obtained from the same sequencing run and assembly rather than from separate virome libraries. The six studies were sequenced on Illumina HiSeq or NovaSeq platforms at raw depths of 25.4–59.4 Gbp (a 2.3-fold range; Table S12). For each study and function, viral contribution was calculated as the proportion of viral gene counts relative to the total number of annotated genes for that function in the same metagenome. Because this proportion was calculated within each sample, sequencing depth was not used to normalize the ratio. These six selected comparisons were analyzed descriptively because only one to seven viral gene calls were identified in each; they were not used to estimate a population-wide mean viral contribution. The targeted functions were chitinase (K03791/PF00182), ammonia assimilation (K01915/PF00120), nitrogen fixation (K02588/PF00142), antibiotic inactivation (K00662/PF02522), and antibiotic inactivation with target alteration (K03587/PF03717), following the subcategory assignments of MetaFuncDecoder v1.0.0. Full study metadata—sequencing platform, raw depth, geographic location, soil type, and management history—are reported in Tables S11 and S12.

### Software, statistics and reproducibility

Functional category assignment was performed with MetaFuncDecoder v1.0.0, a Python tool developed for this study (DOI: 10.5281/zenodo.19009291). All downstream analyses, statistics, and figures were produced in R v4.1.2 using data.table v1.14.8, tidyverse v2.0.0, ggplot2 v3.4.2, patchwork v1.3.0, gridExtra v2.3, RColorBrewer v1.1-3, and openxlsx v4.2.5.2. Group proportions were compared with Fisher’s exact or χ² tests as indicated for each analysis; all tests were two-sided, and exact binomial (Clopper-Pearson) confidence intervals were used for proportions. Analysis scripts were numbered in execution order and were mapped to specific reported analyses; all scripts are archived as described under Code availability.

## Supporting information

All Supplementary tables and figures

## Data availability

A reanalysis of the GSV Atlas described by Graham et al. (2) was performed; no new sequencing data were generated. The GSV Atlas is available for download at https://doi.org/10.25584/2229733. File 2 (data associated with each contig passing QA/QC; 49,649 contigs), File 4 (data associated with each gene; 1,432,147 genes), and File 5 (geographic and physicochemical data of the curated soil samples; 2,953 samples) were used. The six matched total metagenomes used in the viral-versus-total comparison were retrieved from the JGI Integrated Microbial Genomes and Microbiomes (IMG/M) platform on 10 January 2026 under IMG Taxon Object IDs 3300005471, 3300025711, 3300025925, 3300036781, 3300038415, and 3300038418; the corresponding assembly README files were downloaded from the JGI Genome Portal on 8 June 2026 (download ID 1977383). Supplemental Tables S1–S16 are provided with this article, in which the functional annotation dataset is decomposed by database support, evidence specificity, ecosystem, and lysis-cassette status. The complete per-gene annotation table underlying these summaries (metagenome_confidence_annotations.csv; 5,760 MetaFuncDecoder calls, including broad and lysis-associated annotations excluded from the main analysis) is deposited on figshare (doi.org/10.6084/m9.figshare.33780058).

## Code availability

MetaFuncDecoder v1.0.0 is available at github.com/dominikmerges/metafuncdecode and is archived at https://doi.org/10.5281/zenodo.19009291. All analysis, statistical, and figure-generation code associated with this publication is available at github.com/dominikmerges/soil-viral-amg-reanalysis. The numbered analysis scripts, a run_all.R driver by which every reported result can be reproduced from the downloaded GSV Atlas files, the derived result tables and figures, and sessionInfo() output for each script are included in the repository.

## Acknowledgements

I thank the editor and two anonymous reviewers for their constructive comments, which substantially improved the manuscript.

