## Supplementary material for "Predominance of viral core functions over auxiliary metabolism among carbohydrate-active enzymes annotated in the global soil virosphere": All Supplementary tables and figures

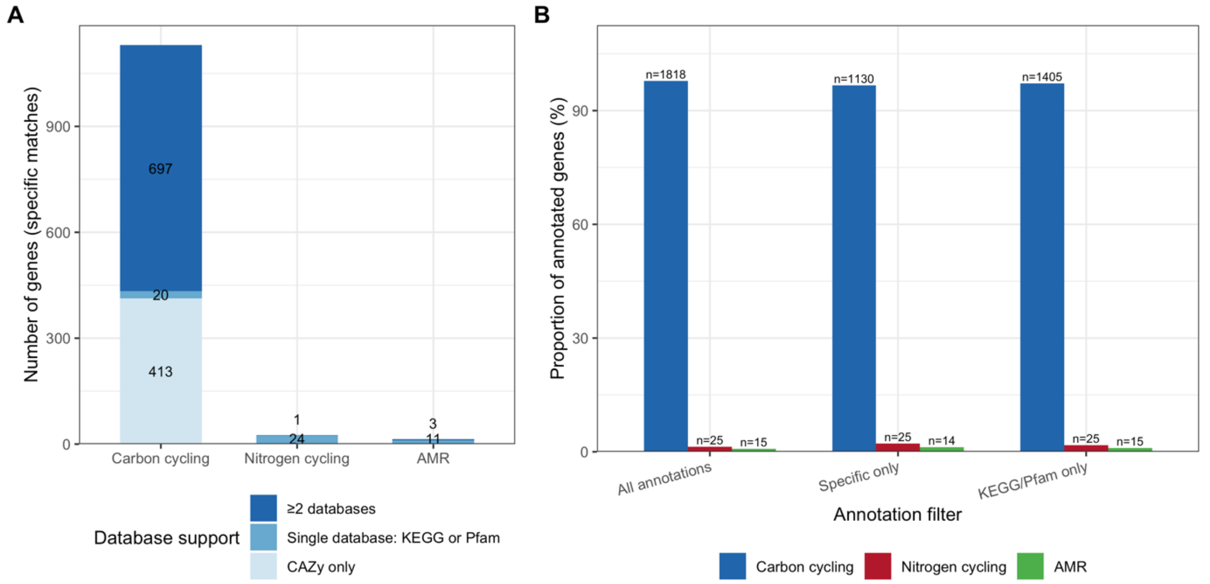


**Figure S1.** Database support and sensitivity to annotation filtering.. Peptidoglycanase (lysis) genes were excluded. (A) Database support for specific (enzyme-resolved) annotations per functional category. MetaFuncDecoder integrates CAZy, KEGG, and Pfam; genes are stratified by whether they were supported by ≥2 databases, by a single database (KEGG or Pfam), or by CAZy alone. Nitrogen-cycling and AMR categories have no CAZy component by definition. Numbers within each bar segment give the count of genes with that level of database support; for the short Nitrogen-cycling and AMR bars, the small ≥2-database count is printed just above the bar. (B) Proportion of annotated genes per functional category under three filters: all annotations, specific matches only (subcategory assigned by MetaFuncDecoder), and KEGG/Pfam-supported only (CAZy-only hits removed). Gene counts are shown above each bar. Carbon-cycling dominance persists across all filters.

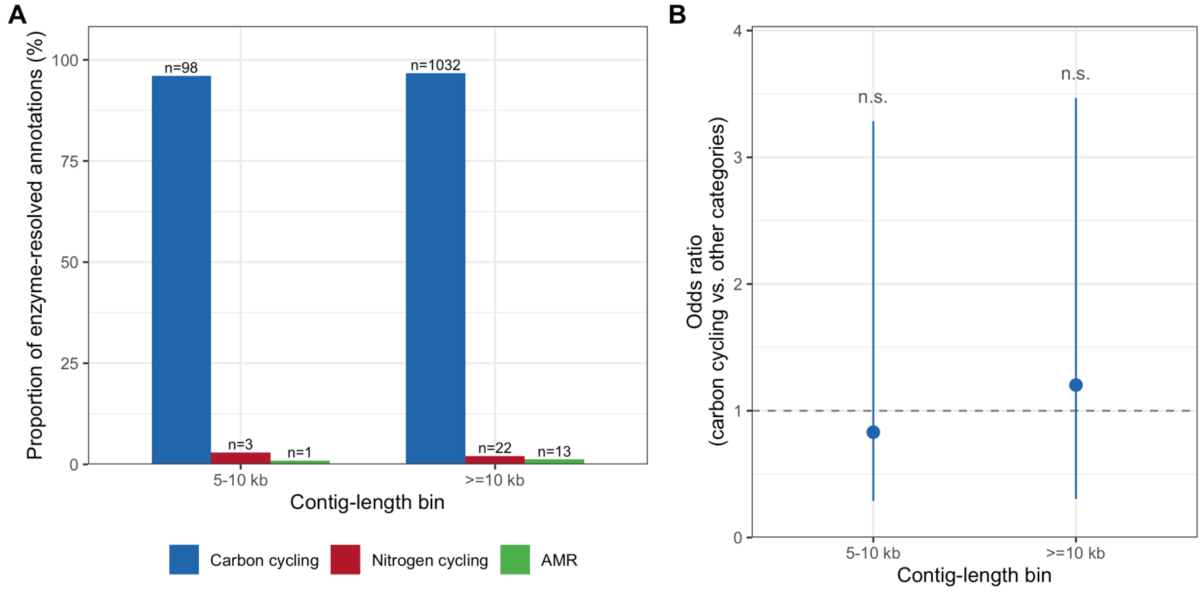


**Figure S2.** Contig-length stratification of viral AMG functional annotations. Peptidoglycanase (lysis) genes were excluded. (A) Proportion of annotated genes assigned to each functional category (carbon cycling, nitrogen cycling, and antimicrobial resistance) within the 5–10 kb and ≥10 kb contig-length bins. Gene counts are shown above each bar. The 1–5 kb bin (n = 80 contigs, 365 genes) contained no functional annotations and is omitted. (B) Odds ratio (Fisher's exact test) for carbon-cycling enrichment relative to all other functional categories, compared between bins. Error bars indicate 95% confidence intervals; the dashed line marks OR = 1 (no enrichment difference). n.s., not significant ($p = 0.77$). Both the 5–10 kb and ≥10 kb contig classes were evaluated; the 1–5 kb class contained no targeted functional annotations.

**Table S1.** Database decomposition of functional annotations by evidence specificity.

| Functional category | Evidence type | No. genes | CAZy | KEGG | Pfam | CAZy only | KEGG or Pfam | Multi-database | % 3 databases | % 2 databases | % 1 database |
|---|---|---|---|---|---|---|---|---|---|---|---|
| | Specific | 14 | 0 | 10 | 7 | 0 | 14 | 3 | 0 | 21.4 | 78.6 |
| *Antibiotics* | Broad | 1 | 0 | 0 | 1 | 0 | 1 | 0 | 0 | 0 | 100 |
| | Specific | 1130 | 1102 | 365 | 557 | 413 | 717 | 697 | 17.4 | 44.2 | 38.3 |
| *Carbon Cycling* | Broad | 688 | 0 | 0 | 688 | 0 | 688 | 0 | 0 | 0 | 100 |
| *Nitrogen Cycling* | Specific | 25 | 0 | 10 | 16 | 0 | 25 | 1 | 0 | 4 | 96 |

**Table S2.** Normalized AMG rates per ecosystem. Raw AMG counts and rates per 10,000 viral genes, stratified by functional category. Peptidoglycanase and broad (non-enzyme-resolved) matches excluded. Normalization denominator is the total number of viral genes (all genes on geNomad-identified viral contigs) per ecosystem.

| Ecosystem | Total samples | Samples with viral genes | Total viral genes | Carbon cycling AMGs | Nitrogen cycling AMGs | Antibiotic resistance AMGs | Total AMGs | Carbon per 10k viral genes | Nitrogen per 10k viral genes | Antibiotic res. per 10k viral genes | Total AMGs per 10k viral genes |
|---|---|---|---|---|---|---|---|---|---|---|---|
| Volcanic | 17 | 15 | 6232 | 7 | 0 | 0 | 7 | 11.23 | 0.00 | 0.00 | 11.23 |
| Plant-associated | 114 | 111 | 253193 | 257 | 0 | 2 | 259 | 10.15 | 0.00 | 0.08 | 10.23 |
| Desert | 31 | 13 | 2001 | 2 | 0 | 0 | 2 | 10.00 | 0.00 | 0.00 | 10.00 |
| Permafrost | 108 | 26 | 19438 | 18 | 1 | 0 | 19 | 9.26 | 0.51 | 0.00 | 9.77 |
| Forest | 528 | 207 | 219008 | 208 | 1 | 3 | 212 | 9.50 | 0.05 | 0.14 | 9.68 |
| Grassland | 119 | 55 | 139105 | 125 | 6 | 1 | 132 | 8.99 | 0.43 | 0.07 | 9.49 |
| Tropical rainforest | 12 | 12 | 3179 | 3 | 0 | 0 | 3 | 9.44 | 0.00 | 0.00 | 9.44 |
| Agricultural | 593 | 153 | 353480 | 284 | 15 | 3 | 302 | 8.03 | 0.42 | 0.08 | 8.54 |
| Unclassified | 1012 | 270 | 49928 | 37 | 0 | 2 | 39 | 7.41 | 0.00 | 0.40 | 7.81 |
| Peat | 398 | 349 | 384077 | 189 | 2 | 3 | 194 | 4.92 | 0.05 | 0.08 | 5.05 |
| Salt marsh | 14 | 8 | 2048 | 0 | 0 | 0 | 0 | 0.00 | 0.00 | 0.00 | 0.00 |
| Shrubland | 7 | 4 | 458 | 0 | 0 | 0 | 0 | 0.00 | 0.00 | 0.00 | 0.00 |

**Table S3.** Subcategory breakdown of specific functional annotations by database support.

| Functional category | Subcategory | No. genes | CAZy | KEGG | Pfam | Multi-database | Three database support | Two database support | One database support |
|---|---|---|---|---|---|---|---|---|---|
| | Antibiotic Efflux | 5 | 0 | 2 | 3 | 0 | 0 | 0 | 5 |
| | Antibiotic Target Alteration | 5 | 0 | 5 | 0 | 0 | 0 | 0 | 5 |
| | Antibiotic Inactivation | 3 | 0 | 2 | 3 | 2 | 0 | 2 | 1 |
| *Antibiotics* | Antibiotic Inactivation / Antibiotic Target Alteration | 1 | 0 | 1 | 1 | 1 | 0 | 1 | 0 |
| | Chitinase | 487 | 471 | 340 | 256 | 388 | 192 | 196 | 99 |
| | β-1,3-Glucanase (GH16) | 133 | 133 | 1 | 113 | 113 | 1 | 112 | 20 |
| | Pectinase | 111 | 111 | 0 | 4 | 4 | 0 | 4 | 107 |
| | Mannanase | 84 | 84 | 0 | 67 | 67 | 0 | 67 | 17 |
| | Other Hemicellulase | 80 | 72 | 9 | 35 | 35 | 1 | 34 | 45 |
| | Arabinogalactanase / β-Glucosidase / Cellulase / Mannanase / Xylanase | 46 | 46 | 0 | 34 | 34 | 0 | 34 | 12 |
| | Alginate/polysaccharide lyase | 42 | 42 | 0 | 0 | 0 | 0 | 0 | 42 |
| | β-Glucosidase | 41 | 41 | 0 | 9 | 9 | 0 | 9 | 32 |
| | Cellulase | 36 | 36 | 0 | 31 | 31 | 0 | 31 | 5 |
| | Xylanase | 17 | 16 | 4 | 5 | 5 | 3 | 2 | 12 |
| | β-1,3-Glucanase (other GH) | 16 | 16 | 4 | 0 | 4 | 0 | 4 | 12 |
| | Arabinofuranosidase / Xylanase | 9 | 9 | 0 | 0 | 0 | 0 | 0 | 9 |
| | Lignin Degradation | 8 | 8 | 0 | 0 | 0 | 0 | 0 | 8 |
| | Starch Degradation | 8 | 5 | 3 | 2 | 2 | 0 | 2 | 6 |
| | Arabinofuranosidase | 7 | 7 | 4 | 0 | 4 | 0 | 4 | 3 |
| *Carbon Cycling* | Arabinogalactanase | 5 | 5 | 0 | 1 | 1 | 0 | 1 | 4 |
| | Nitrogen Fixation | 15 | 0 | 1 | 14 | 0 | 0 | 0 | 15 |
| | Denitrification | 8 | 0 | 8 | 0 | 0 | 0 | 0 | 8 |
| *Nitrogen Cycling* | Ammonia Assimilation | 2 | 0 | 1 | 2 | 1 | 0 | 1 | 1 |

**Table S4.** Subcategory breakdown for Figures 1B-D. Gene counts per enzyme group / mechanism within each functional category. Only specific (enzyme-resolved) annotations included.

| Category | Subcategory | Gene count | % |
|---|---|---|---|
| Carbon cycling | Chitinase | 487 | 43.1 |
| Carbon cycling | Hemicellulase | 197 | 17.4 |
| Carbon cycling | β-1,3-glucanase | 149 | 13.2 |
| Carbon cycling | Cellulase | 123 | 10.9 |
| Carbon cycling | Pectinase | 111 | 9.8 |
| Carbon cycling | Polysaccharide lyase | 42 | 3.7 |
| Carbon cycling | Other | 21 | 1.9 |
| Nitrogen cycling | Nitrogen fixation | 15 | 60 |
| Nitrogen cycling | Denitrification | 8 | 32 |
| Nitrogen cycling | Ammonia assimilation | 2 | 8 |
| Antibiotic resistance | Efflux | 5 | 35.7 |
| Antibiotic resistance | Target alteration | 5 | 35.7 |
| Antibiotic resistance | Inactivation | 3 | 21.4 |
| Antibiotic resistance | Inactivation + Target alteration | 1 | 7.1 |

**Table S5.** Lysis-filter effect on the enzyme-resolved carbon-cycling annotation set. Gene counts per enzyme group before and after removing peptidoglycanase (lysis) genes, with each group's share of the unfiltered carbon-cycling set (n = 5,032) and of the filtered carbon-AMG set (n = 1,130).

| enzyme_group | N in unfiltered carbon-cycling set | N in filtered carbon-AMG set | % of unfiltered set | % of filtered set |
|---|---|---|---|---|
| Peptidoglycanase (lysis) | 3902 | | 77.54372019 | |
| Chitinase | 487 | 487 | 9.678060413 | 43.09734513 |
| Hemicellulase | 197 | 197 | 3.914944356 | 17.43362832 |
| β-1,3-glucanase | 149 | 149 | 2.96104928 | 13.18584071 |
| Cellulase | 123 | 123 | 2.444356121 | 10.88495575 |
| Pectinase | 111 | 111 | 2.205882353 | 9.82300885 |
| Polysaccharide lyase | 42 | 42 | 0.83465819 | 3.71681416 |
| Other | 21 | 21 | 0.41732909 | 1.85840708 |

**Table S6.** CAZyme families in the filtered AMG set. For each enzyme group, the assigned CAZy family, its curated family class (documented virion-structural/lysis role vs. polysaccharide-degrading with no reported viral structural role), and gene count (N).

| enzyme_group | cazy_fam | Curated family class | N |
|---|---|---|---|
| Chitinase | GH19 | Documented virion-structural or lysis role | 381 |
| β-1,3-glucanase | GH16 | Polysaccharide-degrading, no viral structural role reported | 133 |
| Hemicellulase | GH26 | Polysaccharide-degrading, no viral structural role reported | 47 |
| Cellulase | GH5 | Polysaccharide-degrading, no viral structural role reported | 46 |
| Cellulase | GH39 | Polysaccharide-degrading, no viral structural role reported | 39 |
| Chitinase | CBM32 | Non-catalytic binding module | 37 |
| Polysaccharide lyase | PL7 | Documented virion-structural or lysis role | 33 |
| Hemicellulase | CE3 | Polysaccharide-degrading, no viral structural role reported | 29 |
| Pectinase | PL9 | Documented virion-structural or lysis role | 28 |
| Hemicellulase | CE4 | Polysaccharide-degrading, no viral structural role reported | 26 |
| Pectinase | GH140 | Polysaccharide-degrading, no viral structural role reported | 25 |
| Pectinase | PL1_2 | Documented virion-structural or lysis role | 22 |
| Hemicellulase | GH5_40 | Polysaccharide-degrading, no viral structural role reported | 19 |
| Chitinase | GH46 | Polysaccharide-degrading, no viral structural role reported | 17 |
| Cellulase | GH12 | Polysaccharide-degrading, no viral structural role reported | 17 |
| Chitinase | (no CAZy family) | No CAZy family (KEGG/Pfam only) | 16 |
| Chitinase | GH18 | Polysaccharide-degrading, no viral structural role reported | 14 |
| Chitinase | CBM50 | Non-catalytic binding module | 14 |
| Pectinase | CBM67 | Non-catalytic binding module | 10 |
| Pectinase | GH28 | Documented virion-structural or lysis role | 10 |
| Hemicellulase | CE1 | Polysaccharide-degrading, no viral structural role reported | 10 |
| β-1,3-glucanase | GH128 | Polysaccharide-degrading, no viral structural role reported | 9 |
| Hemicellulase | (no CAZy family) | No CAZy family (KEGG/Pfam only) | 9 |
| Cellulase | GH5_5 | Polysaccharide-degrading, no viral structural role reported | 9 |
| Pectinase | GH145 | Polysaccharide-degrading, no viral structural role reported | 7 |
| Hemicellulase | GH141 | Polysaccharide-degrading, no viral structural role reported | 7 |
| Hemicellulase | CBM35 | Non-catalytic binding module | 7 |
| Hemicellulase | GH5_7 | Polysaccharide-degrading, no viral structural role reported | 7 |
| Chitinase | GH75 | Polysaccharide-degrading, no viral structural role reported | 7 |
| Hemicellulase | CBM13 | Non-catalytic binding module | 6 |
| Polysaccharide lyase | PL6 | Documented virion-structural or lysis role | 5 |
| Hemicellulase | GH54 | Polysaccharide-degrading, no viral structural role reported | 5 |
| Hemicellulase | GH10 | Polysaccharide-degrading, no viral structural role reported | 4 |
| Hemicellulase | CE6 | Polysaccharide-degrading, no viral structural role reported | 4 |
| Hemicellulase | GH30 | Polysaccharide-degrading, no viral structural role reported | 4 |
| Polysaccharide lyase | PL14 | Documented virion-structural or lysis role | 4 |
| β-1,3-glucanase | GH81 | Polysaccharide-degrading, no viral structural role reported | 4 |
| Other (Starch degradation) | GH87 | Polysaccharide-degrading, no viral structural role reported | 4 |
| Pectinase | PL1 | Documented virion-structural or lysis role | 4 |
| Pectinase | GH137 | Polysaccharide-degrading, no viral structural role reported | 4 |
| Other (Lignin degradation) | AA7 | Polysaccharide-degrading, no viral structural role reported | 3 |
| Cellulase | GH9 | Polysaccharide-degrading, no viral structural role reported | 3 |

| | | | |
|---|---|---|---|
| Cellulase | GH5_1 | Polysaccharide-degrading, no viral structural role reported | 3 |
| Hemicellulase | GH43 | Polysaccharide-degrading, no viral structural role reported | 3 |
| Other (Lignin degradation) | AA3 | Polysaccharide-degrading, no viral structural role reported | 3 |
| Other (Arabinogalactanase) | GH43_37 | Polysaccharide-degrading, no viral structural role reported | 3 |
| Other (Starch degradation) | (no CAZy family) | No CAZy family (KEGG/Pfam only) | 3 |
| Hemicellulase | GH113 | Polysaccharide-degrading, no viral structural role reported | 3 |
| β-1,3-glucanase | GH64 | Polysaccharide-degrading, no viral structural role reported | 2 |
| Cellulase | GH1 | Polysaccharide-degrading, no viral structural role reported | 2 |
| Other (Lignin degradation) | AA5 | Polysaccharide-degrading, no viral structural role reported | 2 |
| Cellulase | GH5_39 | Polysaccharide-degrading, no viral structural role reported | 2 |
| Other (Starch degradation) | GH32 | Polysaccharide-degrading, no viral structural role reported | 1 |
| Hemicellulase | GH93 | Polysaccharide-degrading, no viral structural role reported | 1 |
| Cellulase | GH5_46 | Polysaccharide-degrading, no viral structural role reported | 1 |
| Hemicellulase | GH5_10 | Polysaccharide-degrading, no viral structural role reported | 1 |
| Hemicellulase | GH27 | Polysaccharide-degrading, no viral structural role reported | 1 |
| Hemicellulase | GH51 | Polysaccharide-degrading, no viral structural role reported | 1 |
| Pectinase | GH78 | Polysaccharide-degrading, no viral structural role reported | 1 |
| β-1,3-glucanase | GH144 | Polysaccharide-degrading, no viral structural role reported | 1 |
| Chitinase | CBM12 | Non-catalytic binding module | 1 |
| Hemicellulase | GH2 | Polysaccharide-degrading, no viral structural role reported | 1 |
| Hemicellulase | CE2 | Polysaccharide-degrading, no viral structural role reported | 1 |
| Other (Arabinogalactanase) | GH43_24 | Polysaccharide-degrading, no viral structural role reported | 1 |
| Hemicellulase | CBM9 | Non-catalytic binding module | 1 |
| Cellulase | GH8 | Polysaccharide-degrading, no viral structural role reported | 1 |
| Other (Arabinogalactanase) | CBM61 | Non-catalytic binding module | 1 |

**Table S7.** Chitinase-associated family breakdown. CAZy families annotated as chitinase-associated, each family's curated class (virion-structural/lysis vs. degradative), gene count (N) and percentage of the chitinase-associated set (pct).

| cazy_fam | family_class | N | pct |
|---|---|---|---|
| GH19 | Documented virion-structural or lysis role | 381 | 78.2 |
| CBM32 | Non-catalytic binding module | 37 | 7.6 |
| GH46 | Polysaccharide-degrading, no viral structural role reported | 17 | 3.5 |
| (no CAZy family) | No CAZy family (KEGG/Pfam only) | 16 | 3.3 |
| GH18 | Polysaccharide-degrading, no viral structural role reported | 14 | 2.9 |
| CBM50 | Non-catalytic binding module | 14 | 2.9 |
| GH75 | Polysaccharide-degrading, no viral structural role reported | 7 | 1.4 |
| CBM12 | Non-catalytic binding module | 1 | 0.2 |

**Table S8.** Cassette-adjacency of GH19 and control CAZyme families on holin/spanin-bearing viral contigs. Each row is one CAZyme set, plus a random-gene control drawn from the same contigs. n_genes, total genes in the set across all viral contigs; n_with_cassette, the subset located on contigs carrying at least one detectable holin or spanin ORF (the genes eligible for the adjacency test); n_adjacent, of those eligible genes, the number lying within three ORFs of the nearest holin or spanin; pct_adjacent (95% CI), n_adjacent / n_with_cassette with an exact binomial (Clopper–Pearson) interval (binom.test in R); median_dist, median ORF distance to the nearest cassette component among eligible genes. Holin, spanin, peptidoglycan-hydrolase and CAZy assignments are the pre-existing GSVA/geNomad per-gene annotations (Methods), not re-annotated here; only 68 of 381 GH19 genes (17.8%) sit on a contig with a detectable cassette marker and therefore enter this test. These 68 genes sit on 68 distinct contigs, which are also the eligibility set for the random-gene control in Table S9.

| set | n_genes | n_with_cassette | n_adjacent | pct_adjacent | ci_lo | ci_hi | median_dist |
|---|---|---|---|---|---|---|---|
| GH19 chitinase-called | 381 | 68 | 59 | 86.76470588 | 76.35644457 | 93.76516859 | 1 |
| GH24 lysozyme | 1372 | 172 | 146 | 84.88372093 | 78.64079863 | 89.88172184 | 1 |
| GH108 endolysin | 436 | 41 | 37 | 90.24390244 | 76.86854507 | 97.27725332 | 1 |
| GH25 endolysin | 853 | 53 | 34 | 64.1509434 | 49.8033224 | 76.85690531 | 3 |
| GH16 β-1,3-glucanase | 133 | 10 | 3 | 30 | 6.673951118 | 65.24528501 | 33 |
| Random gene same contigs | 380 | 66 | 12 | 18.18181818 | 9.763520202 | 29.60654558 | 12 |

**Table S9.** Fisher tests of lysis-cassette adjacency. Denominators differ slightly by set: n_with_cassette is 68 for GH19 and 66 for the random-gene control. Both sets are evaluated on the same 68 GH19-bearing contigs that carry a detectable holin or spanin ORF. Each GH19 gene on one of these contigs enters the test, giving 68 (one per contig, as no cassette-bearing contig carries more than one GH19 gene). The control draws one non-GH19 gene per contig; on two of these contigs the drawn gene was itself the contig's only holin or spanin ORF, and because a gene cannot be counted as adjacent to itself (the query ORF is removed before the nearest-cassette distance is taken) it has no partner marker and drops out, giving 66. This two-gene difference does not affect the contrast (control 12/66 = 18.2%, or 12/68 = 17.6%, versus 59/68 = 86.8% for GH19). Odds ratios express the odds of adjacency for GH19 relative to each control set.

| comparison | odds_ratio | p_value |
|---|---|---|
| GH19 vs GH24 lysozyme | 1.16671833 | 0.84004049 |
| GH19 vs GH108 endolysin | 0.710871265 | 0.763080357 |
| GH19 vs GH25 endolysin | 3.621999796 | 0.004591589 |
| GH19 vs GH16 β-1,3-glucanase | 14.44700849 | 0.000363752 |
| GH19 vs Random gene same contigs | 28.37136587 | 3.69503E-16 |

**Table S10.** Length-stratified peptidoglycan-hydrolase (PGH) content of GH19-bearing contigs versus all viral contigs. For each length bin: number of contigs, number carrying a separate PGH (a PGH other than the focal GH19), and the percentage with a 95% exact binomial (Clopper–Pearson) confidence interval.

| Length bin (kb) | Contig set | n contigs | n with separate PGH | % (95% CI) |
|---|---|---|---|---|
| <10 | GH19-bearing | 38 | 0 | 0 (0–9.3) |
| <10 | All viral | 18,305 | 1,271 | 6.9 (6.6–7.3) |
| 10–20 | GH19-bearing | 87 | 13 | 14.9 (8.2–24.2) |
| 10–20 | All viral | 16,017 | 2,289 | 14.3 (13.8–14.8) |
| 20–30 | GH19-bearing | 30 | 5 | 16.7 (5.6–34.7) |
| 20–30 | All viral | 5,772 | 1,317 | 22.8 (21.7–23.9) |
| 30–50 | GH19-bearing | 116 | 22 | 19.0 (12.3–27.3) |
| 30–50 | All viral | 6,496 | 2,487 | 38.3 (37.1–39.5) |
| ≥50 | GH19-bearing | 109 | 23 | 21.1 (13.9–30.0) |
| ≥50 | All viral | 3,059 | 1,386 | 45.3 (43.5–47.1) |

**Table S11.** Metadata for the six matched JGI studies used in the viral-versus-total metagenomic comparison.

| JGI Study ID | Study # | Function | Target IDs (KO/Pfam) | Ecosystem | Soil type | Geographic location | Latitude | Longitude | Management history |
|---|---|---|---|---|---|---|---|---|---|
| 3300005471 | S1 | Antibiotic inactivation | K00662/PF02522 | Agricultural | Loam (rhizosphere) | USA: Michigan: Kellogg Biological Station | 42.3948 | -85.3738 | LTER; corn-switchgrass-miscanthus cropping system |

| | | | | | | | | | |
|---|---|---|---|---|---|---|---|---|---|
| 3300025711 | S4 | Chitinase | K03791/PF00182 | Plant-associated | Rhizosphere soil | Michigan, USA | 42.3948 | -85.3738 | LTER; switchgrass cropping system |
| 3300025925 | S3 | Chitinase | K03791/PF00182 | Plant-associated | Rhizosphere soil | USA: Michigan, Kellogg Biological Station | 42.3948 | -85.3738 | LTER; switchgrass cropping system |
| 3300036781 | S2 | Antibiotic inactivation + target alteration | K03587/PF03717 | Forest | Forest soil | USA: California | 37 | -122.2 | Native pine forest; no agricultural management |
| 3300038415 | S5 | Ammonia assimilation | K01915/PF00120 | Peat | Peat | USA: Minnesota | 47.5056 | -93.4534 | Long-term peat bog monitoring (SPRUCE experiment) |
| 3300038418 | S6 | Nitrogen fixation | K02588/PF00142 | Peat | Peat | USA: Minnesota | 47.5056 | -93.4534 | Long-term peat bog monitoring (SPRUCE experiment) |

**Table S12.** Sequencing statistics for the six matched JGI studies used in the viral-versus-total metagenomic comparison. Raw Gbp for HiSeq studies estimated as raw reads x 151 bp (interleaved paired-end fastq). NovaSeq studies report raw base counts directly from JGI README files. 'Same assembly' indicates that viral contigs were identified within the same assembled metagenome as the total metagenomic annotations; viral and total annotations derive from the same sequencing run, not from separate virome libraries. LTER = Long-Term Ecological Research site (Kellogg Biological Station). SPRUCE = Spruce and Peatland Responses Under Changing Environments.

| JGI Study ID | Sequencing platform | Raw reads (M) | Raw Gbp | Gbp source | Filtered reads (M) | Assembler | Assembly date | Assembled size (Gbp) | Gene count | Total KEGG (target) | Viral KEGG (target) | Viral % | Same assembly | Sample name |
|---|---|---|---|---|---|---|---|---|---|---|---|---|---|---|
| 3300005471 | Illumina HiSeq | 375.3 | 56.7 | estimated | 356 | megahit | 2015-06-07 | 4.93 | 12466515 | 322 | 1 | 0.31 | Yes | Corn, switchgrass and miscanthus rhizosphere microbial communities from Kellogg Biological Station, Michigan, USA - KBS K1-50-2 metaG |
| 3300025711 | Illumina HiSeq-2500 | 372.1 | 56.2 | estimated | 372.1 | metaSPAdes 3.10 (IMG) | 2018-03-21 | 0.49 | 1002376 | 71 | 7 | 9.86 | Yes | Switchgrass rhizosphere microbial communities from Kellogg Biological Station, Michigan, USA - KBS S5-4 metaG (SPAdes) |

| | | | | | | | | | | | | | | |
|---|---|---|---|---|---|---|---|---|---|---|---|---|---|---|
| 3300025925 | Illumina HiSeq-2500 | 353.6 | 53.4 | estimated | 353.6 | metaSPAdes 3.10 (IMG) | 2018-03-21 | 3.41 | 7835490 | 95 | 2 | 2.11 | Yes | Switchgrass rhizosphere microbial communities from Kellogg Biological Station, Michigan, USA - KBS S6-3 metaG (SPAdes) |
| 3300036781 | Illumina NovaSeq | 393.2 | 59.4 | reported | 390.5 | metaSPAdes 3.13 | 2019-06-26 | 2.71 | 4215572 | 973 | 1 | 0.1 | Yes | Soil fungal communities from native Pine forests in California, United States - CAS_2 |
| 3300038415 | Illumina NovaSeq S4 | 168 | 25.4 | reported | 162.8 | metaSPAdes 3.13 | 2020-02-27 | 0.69 | 976939 | 366 | 1 | 0.27 | Yes | Peat soil microbial communities from Marcell Experimental Forest, MN, USA - P7_D10 |
| 3300038418 | Illumina NovaSeq S4 | 209.3 | 31.6 | reported | 206.5 | metaSPAdes 3.13 | 2020-02-27 | 0.68 | 925279 | 34 | 1 | 2.94 | Yes | Peat soil microbial communities from Marcell Experimental Forest, MN, USA - P4_D8 |

**Table S13.** Ecosystem sampling overview. Number of metagenome samples, viral gene recovery, zero-gene sample fraction, and broad functional annotation rate (any Pfam/KEGG/CAZy hit) per ecosystem category. 'Unclassified' comprises samples for which GOLD.Specific.Ecosystem was not curated in the IMG/M database. Ecosystem labels were harmonized from GOLD ontology (e.g., 'Forest Soil'/'Forest soil' merged as 'Forest'; 'Agricultural land'/'Agricultural'/'Agricultural soil' merged as 'Agricultural'). Shrubland (n=7, all peptidoglycanase, zero AMGs) is a named GOLD ecosystem pooled into 'Other' by the <30-sample rule. 'Genes with any functional annotation' counts genes matching at least one of Pfam, KEGG or CAZy, independent of the three AMG categories reported in the 'Normalized AMG rates' sheet; this is the broad annotation rate behind the catalogue-wide dark-matter estimate in the Discussion. Ecosystems with fewer than 100 samples (Volcanic, Tropical rainforest, Salt marsh, Desert) rest on 2,001-6,232 genes and their annotation rate should be read with caution.

| Ecosystem | Total samples | Samples with viral genes | Samples without viral genes | % with viral genes | Total viral genes | Mean viral genes per sample | Median viral genes per sample | Genes with any functional annotation (Pfam/KEGG/CAZy) | % annotated (any hit) |
|---|---|---|---|---|---|---|---|---|---|
| Volcanic | 17 | 15 | 2 | 88.2 | 6232 | 366.6 | 143 | 1331 | 21.4 |
| Plant-associated | 114 | 111 | 3 | 97.4 | 253193 | 2221 | 1134.5 | 47758 | 18.9 |
| Desert | 31 | 13 | 18 | 41.9 | 2001 | 64.5 | 0 | 571 | 28.5 |
| Permafrost | 108 | 26 | 82 | 24.1 | 19438 | 180 | 0 | 3611 | 18.6 |
| Forest | 528 | 207 | 321 | 39.2 | 219008 | 414.8 | 0 | 41847 | 19.1 |
| Grassland | 119 | 55 | 64 | 46.2 | 139105 | 1168.9 | 0 | 24558 | 17.7 |
| Tropical rainforest | 12 | 12 | 0 | 100 | 3179 | 264.9 | 116.5 | 727 | 22.9 |
| Agricultural | 593 | 153 | 440 | 25.8 | 353480 | 596.1 | 0 | 61411 | 17.4 |
| Unclassified | 1012 | 270 | 742 | 26.7 | 49928 | 49.3 | 0 | 9035 | 18.1 |
| Peat | 398 | 349 | 49 | 87.7 | 384077 | 965 | 462.5 | 68913 | 17.9 |
| Salt marsh | 14 | 8 | 6 | 57.1 | 2048 | 146.3 | 18.5 | 377 | 18.4 |
| Shrubland | 7 | 4 | 3 | 57.1 | 458 | 65.4 | 17 | 119 | 26 |

**Table S14.** Number of supporting databases per functional annotation, stratified by functional category.

| Functional category | No. genes | 1 database | 2 databases | 3 databases | % 1 database | % 2 databases | % 3 databases | Max. achievable |
|---|---|---|---|---|---|---|---|---|
| *Carbon Cycling* | 1818 | 1121 | 500 | 197 | 61.7 | 27.5 | 10.8 | 3 |
| *Antibiotics* | 15 | 12 | 3 | 0 | 80 | 20 | 0 | 2 |
| *Nitrogen Cycling* | 25 | 24 | 1 | 0 | 96 | 4 | 0 | 2 |

**Table S15.** Identifier sets used to flag lysis-cassette components and to classify CAZy families. Every viral gene was flagged using its pre-existing per-gene Pfam, CAZy and KEGG annotations (Graham et al. 2024, File 4); genes were not re-annotated. Two nested peptidoglycanase definitions are used. The exclusion set (first row — the MetaFuncDecoder peptidoglycanase subcategory) removes glycoside-hydrolase endolysins from the carbon inventory. The endolysin scan (first four rows — the CAZy, glycosidase-Pfam, amidase/endopeptidase-Pfam and KEGG identifiers) is the catalogue-wide flag used for the lysis-cassette co-occurrence test; it is a superset of the exclusion set, adding the amidase and endopeptidase endolysin classes. Holin and spanin sets complete the cassette; the non-catalytic CBM and dual-use CAZy sets are used in the carbon-family classification.

| Category | n | Identifiers | Basis / source |
|---|---|---|---|
| Peptidoglycanase CAZy — glycoside-hydrolase endolysins (MFD exclusion set) | 11 | GH22, GH23, GH24, GH25, GH73, GH102, GH103, GH104, GH108, GH153, CE9 (+ text patterns: lysozyme, muramidase, N-acetylmuramidase, peptidoglycan hydrolase, lytic transglycosylase) | The MetaFuncDecoder peptidoglycanase subcategory (López-Mondéjar et al. 2022, Table S3). Excluded from the carbon inventory AND forms the CAZy part of the endolysin scan. Only GH23/24/25/73/103/104/108 carry genes here (3,652); GH22, GH102, GH153 and CE9 = 0 genes |
| Endolysin Pfam — glycosidase class | 6 | PF00959 (Phage_lysozyme), PF01183 (Glyco_hydro_25), PF05838 (Glyco_hydro_108), PF01464 (SLT / lytic transglycosylase), PF01832 (Glucosaminidase), PF00062 (C-type lysozyme) | Cleave the glycan backbone; overlap the CAZy glycoside-hydrolase families above (Vollmer et al. 2008; Broendum et al. 2018) |
| Endolysin Pfam — amidase / endopeptidase class | 8 | PF01510 (Amidase_2), PF01520 (Amidase_3), PF05257 (CHAP), PF00877 (NlpC/P60), PF01551 (Peptidase_M23), PF08291 (Peptidase_M15_3), PF13539 (Peptidase_M15_4), PF02557 (VanY) | Cleave peptide/amide bonds, not the glycan backbone; not glycoside hydrolases and outside CAZy. Recovered by the endolysin scan but NOT by the exclusion set — 4,249 such amidase/endopeptidase-only endolysins here (Vollmer et al. 2008; Broendum et al. 2018) |
| Endolysin — KEGG | 9 | K01185, K07273, K02395, K03642, K12054, K08309, K08305, K21508, K01448 | Lysozyme, peptidoglycan-hydrolase and lytic-transglycosylase orthologues |
| Holin — Pfam | 36 | PF04020, PF04531, PF04550, PF04688, PF04971, PF05102, PF05105, PF05106, PF05449, PF06946, PF07332, PF09682, PF10746, PF10960, PF11031, PF11351, PF13272, PF16079, PF16080, PF16081, PF16082, PF16083, PF16084, PF16085, PF16931, PF16935, PF16936, PF16938, PF16945, PF23778, PF23809, PF23987, PF24205, PF26844, PF27028, PF28015 | All Pfam families whose entry name denotes a holin or antiholin (InterPro query) |
| Spanin — Pfam | 3 | PF03245, PF17531, PF26831 | Outer-membrane-disruption component of the lysis cassette |
| Non-catalytic carbohydrate-binding module (CBM) | 8 | CBM9, CBM12, CBM13, CBM32, CBM35, CBM50, CBM61, CBM67 | Carbohydrate-binding modules with no hydrolytic activity (CAZy database) |
| Dual-use / virion-structural CAZy | 8 | GH19, PL1, PL1_2, PL6, PL7, PL9, PL14, GH28 | Families with a documented virion-structural, host-entry or lysis role (GH19: Orlando et al. 2021, Edvardsen et al. 2025, Meng et al. 2024; PLs and GH28: Latka et al. 2017, Martin et al. 2025) |

**Table S16.** Ecosystem label harmonization. Mapping of GOLD ontology metadata to the ecosystem categories used in this study.

| Ecosystem bin | Source column | Original labels | Rationale |
|---|---|---|---|
| Agricultural | GOLD.Specific.Ecosystem | Agricultural land; Agricultural; Agricultural soil | Synonym harmonization |
| Forest | GOLD.Specific.Ecosystem | Forest Soil; Forest soil | Case harmonization |
| Grassland | GOLD.Specific.Ecosystem | Grasslands | Direct mapping |
| Permafrost | GOLD.Specific.Ecosystem | Permafrost | Direct mapping |
| Desert | GOLD.Specific.Ecosystem | Desert | Direct mapping |
| Tropical rainforest | GOLD.Specific.Ecosystem | Tropical rainforest | Direct mapping |
| Shrubland | GOLD.Specific.Ecosystem | Shrubland | Named ecosystem, <30 samples -> pooled as Other |
| Peat | GOLD.Ecosystem.Type | Peat (all subtypes) | Distinct ecosystem type in GOLD |
| Plant-associated | GOLD.Ecosystem.Type | Rhizosphere; Roots; Rhizoplane | All plant-associated soil types combined |
| Volcanic | GOLD.Ecosystem.Type | Volcanic | Distinct ecosystem type in GOLD |
| Salt marsh | GOLD.Ecosystem.Type | Marine (all = Salt marsh) | All marine samples were salt marsh |
| Unclassified | Various | Soil/Unclassified; Unclassified/Unclassified; Soil/Soil | Metadata gap in IMG/GOLD database |
